# Zika virus-induced fetal demise is driven by strain- and dose-specific RLR-driven activation of the interferon response in the decidua, placenta, and fetus in *Ifnar1^-/-^* mice

**DOI:** 10.1101/2025.02.12.637947

**Authors:** Ellie K. Bohm, David Castañeda, Qun Lu, Michael D. Cameron, Matthew T. Aliota

## Abstract

Congenital Zika syndrome (CZS), the set of fetal and neonatal complications associated with Zika virus (ZIKV) infection in pregnancy, was first noted during the outbreak in the Americas in 2015-16. However, there was an unequal distribution of ZIKV cases and severe outcomes in all areas where ZIKV emerged in the Americas, demonstrating that the risk of CZS varied over space and time. Recently, we demonstrated that phenotypic heterogeneity existed between closely-related ZIKV strains. All ZIKV strains tested infected the placenta but varied in their capacity to cause overt fetal harm. Here, we further characterized the relative contributions of virus genotype and infecting dose of two phenotypically distinct ZIKV strains across multiple timepoints in gestation in pregnant mice that lack type-I interferon receptor function (*Ifnar1^-/-^*). To better understand the underlying causes of adverse fetal outcomes, we used RNA sequencing to compare ZIKV-infected and uninfected tissues. We found that ZIKV infection triggers retinoic acid-inducible gene I (RIG-I)-like receptor-mediated activation of the interferon response at the maternal-fetal interface. However, modest chemical inhibition of RIG-I activation in the decidua and placenta did not protect against fetal demise. Instead, the fetal interferon response was significantly associated with fetal demise. Together, these findings suggest that the response to ZIKV at the maternal-fetal interface can vary depending on the infecting ZIKV genotype and dose, and that the fetal immune response is an important mediator of fetal harm.

**IMPORTANCE:** Previously, we used a mouse model of ZIKV infection during pregnancy to assess the pathogenic potential to the fetus of a panel of five, low-passage ZIKV strains representing the viral genetic diversity in the Americas. We found that phenotypic heterogeneity existed between these closely-related ZIKV strains. Here, we show that this heterogeneity is driven by retinoic acid-inducible gene I (RIG-I)-like receptor-mediated activation of the interferon response at the maternal-fetal interface. We used chemical inhibition of the RIG-I pathway and measured the transcriptional activity of interferon stimulated genes in fetuses to demonstrate that the fetal immune response may contribute to fetal demise.

## INTRODUCTION

Zika virus (ZIKV) infection during pregnancy can cause a spectrum of adverse fetal outcomes collectively termed Congenital Zika Syndrome (CZS), but not all children exposed to ZIKV *in utero* develop these abnormalities. While it is well-established that ZIKV can cause fetal harm, *how* ZIKV causes fetal harm remains unclear. Whether fetal pathology manifests for a given pregnancy is dependent on myriad factors including gestational age of the fetus, maternal immunity, maternal-fetal barrier integrity, and ZIKV tropism (1, 2). ZIKV can be vertically transmitted through the maternal-fetal barrier, but the route and frequency of transmission remains uncertain. It is thought that ZIKV is vertically transmitted from maternal circulation to the maternal-derived decidua, then to the adjacent fetal-derived placenta, and finally to the fetus (3, 4). ZIKV can replicate in several cell types of the human maternal-fetal interface (MFI) including maternal decidual cells (5), fetal trophoblast cells, and fetal endothelial cells (3). Many studies report detection of viral proteins and/or viral RNA (vRNA) in the placental tissues of ZIKV-infected pregnant people (6). Many animal studies recapitulate these findings with ZIKV vRNA detected in multiple MFI tissues of non-human primates (7, 8), but these same studies were unable to determine the route of transmission through the tissues. Human cohort studies report varying frequencies of infection of the MFI and fetus. In one case study, over half of ZIKV-infected mothers had ZIKV vRNA detected in placental and/or fetal tissue at term (9). In cases of severe microcephaly, evidence of fetal infection was relatively common (10–12), indicating that fetal infection is likely one mechanism of fetal harm. But with limited screening of apparently-normal infants who have subtle neurological sequelae, it remains unknown if fetal infection is a precursor in all cases of CZS.

Ultimately, fetal infection may not be required for fetal harm. Recent cohort studies show that infants with CZS have high levels of inflammatory markers (13, 14), suggesting a significant inflammatory response before birth. A robust inflammatory response can cause placental dysfunction, a syndrome during which the placenta fails to develop properly and deliver nutrients, blood, and oxygen to the growing fetus. Placental dysfunction results in intrauterine growth restriction, abnormal development, and miscarriage (15), which have been observed in neonates and infants with CZS. ZIKV vRNA persistence at the MFI can also induce high levels of interferon (IFN) (16). In some animal models, placental damage caused by the IFN response was a precursor to fetal demise, and fetal infection was not required (17–21). Consistent with this, certain nucleotide polymorphisms in IFN receptors and immune profiles were associated with higher levels of IFN-stimulated genes and increased risk of CZS in humans (22, 23). Together, these findings suggest that ZIKV infection of the fetus is not required in all cases of fetal harm.

Epidemiological data from the 2015-2016 American outbreak showed that although Asian/American-lineage ZIKV strains share >99% nucleotide-identity (24), they cause heterogeneous rates of fetal harm (25–31). This suggests ongoing virus evolution during the 2015-2016 outbreak in the Americas may have given rise to phenotypic variants that differ in the mechanism by which developing fetuses are harmed. Indeed, we unexpectedly found that phenotypic heterogeneity existed between closely-related ZIKV strains in a pregnant *Ifnar1^-/-^*mouse model (18). The Asian-lineage ZIKVs we tested had varying capacities to cause fetal demise, ranging between 9 - 51%—importantly, demise occurred in the absence of detectable fetal infection(18). Other infection parameters, including maternal viremia, placental infection, placental histopathology, and intrauterine growth restriction were similarly heterogeneous in our mouse model (18). Surprisingly, none of these phenotypes positively correlated with the rate of fetal demise.

Therefore, to identify other factors that may contribute to ZIKV-induced fetal demise, we leveraged the natural variability in phenotype that exists between closely-related ZIKV strains and initiated transcriptome profiling studies to assess gene expression changes in the placenta. We used two ZIKV strains that showed different pregnancy phenotypes: a strain from Brazil, ZIKV-BRA (Paraiba_01), that causes significant fetal demise, and a strain from Mexico, ZIKV-MEX (R116265), that does not. We found that ZIKV infection results in strain- and dose-dependent activation of the IFN response at the MFI prior to fetal demise. Further analysis suggested that retinoic acid-inducible gene I (RIG-I) sensing of ZIKV vRNA was a primary driver of the IFN response. Since the IFN response is known to be pathogenic during pregnancies (17, 21), we aimed to investigate if chemical inhibition of RIG-I signaling reduced rates of fetal demise following ZIKV-BRA infection. We found that modest RIG-I inhibition at the MFI does not protect against fetal demise, but identified a strong association between an increased fetal IFN response and fetal demise.

## RESULTS

### ZIKV strain- and dose-dependent pregnancy phenotypes are present across gestation

Previously, we determined that there is strain-dependent phenotypic heterogeneity in pregnancy outcomes following *in utero* ZIKV exposure in pregnant *Ifnar1^-/-^* mice (18). We compared a panel of five geographically-distinct, low-passage Asian/American-lineage ZIKV strains and assessed pregnancy outcomes at a single necropsy timepoint (E14.5) to evaluate the extent to which pregnancy outcomes varied by infecting ZIKV genotype. Viruses from Brazil and Cambodia caused significantly more embryo resorption than viruses from Panama, Puerto Rico, and Mexico (18). Now, to determine when strain-dependent outcomes manifest and assess the influence of dose, we compared pregnancy outcomes at multiple points in gestation. We inoculated with 10^3^ PFU ZIKV-MEX, 10^5^ PFU ZIKV-MEX, and 10^3^ PFU ZIKV-BRA. We chose these two ZIKV strains because they have distinct pregnancy phenotypes—ZIKV-BRA causes significant fetal resorption and ZIKV-MEX does not—when inoculated with 10^3^ PFU (18). These virus strains differ by only seven amino acids (**Table 1**). We included a high-dose inoculation of ZIKV-MEX (10^5^ PFU ZIKV-MEX) to determine if increasing the dose for this virus strain impacts the rate of fetal resorption. To assess pregnancy outcomes, *Ifnar1*^-/-^ dams were time-mated with wildtype (WT) males to produce fetal and placental tissue with intact IFN signaling, as we have done previously (18, 19, 32). Pregnant *Ifnar1*^-/-^ dams then were inoculated with 10^3^ or 10^5^ PFU ZIKV-MEX or 10^3^ PFU ZIKV-BRA via subcutaneous footpad inoculation at embryonic day 7.5 (E7.5). E7.5 corresponds to the mid-to-late first trimester in humans (33). Dams were monitored daily for clinical signs until the time of necropsy; no overt clinical signs were observed in any virus- or PBS-inoculated dams. We collected serum at 2, 4, 7 (10^5^ PFU ZIKV-MEX only), and 10 days post inoculation (dpi) to compare maternal viremia kinetics between the two viruses. At 2 dpi, all groups were significantly different from each other (p < 0.0409), with 10^5^ PFU ZIKV-MEX-inoculated animals having the highest serum titers (**Figure 1A**). At 4 dpi, the 10^3^ PFU ZIKV-BRA group had significantly higher titers than both ZIKV-MEX groups (p < 0.0001). At 10 dpi, titers did not differ significantly between 10^3^ PFU ZIKV-MEX, 10^5^ PFU ZIKV-MEX, and 10^3^ PFU ZIKV-BRA, and were largely undetectable. There was no significant difference between 10^5^ PFU ZIKV-MEX and 10^3^ PFU ZIKV-MEX and 10^3^ PFU ZIKV-BRA from reference (Bohm 2021) at 7 dpi (p > 0.9999).

**Figure 1:**
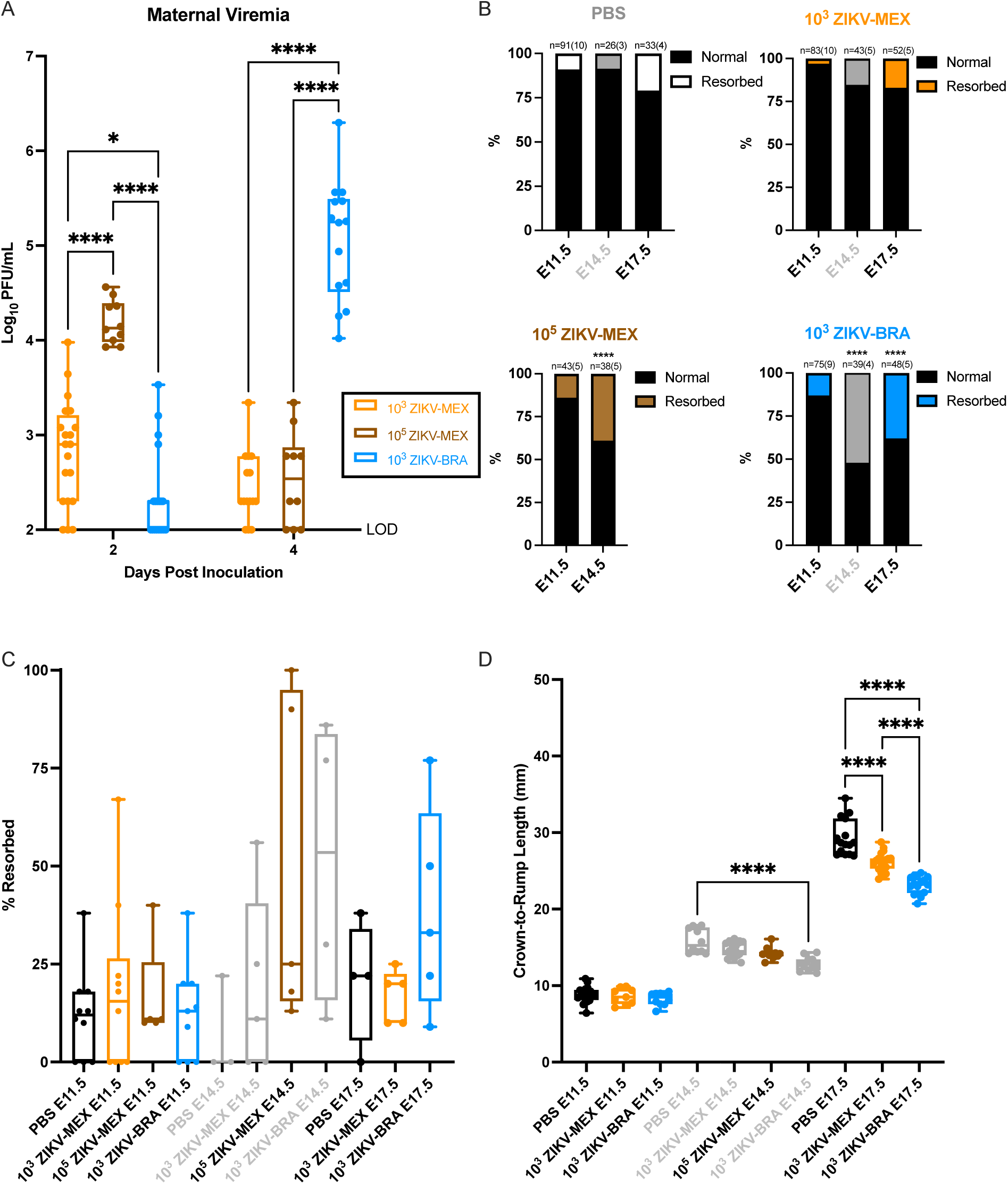
ZIKV strain phenotypic heterogeneity is present across gestation. (A) Time-mated *Ifnar1*^−/−^ dams were inoculated with 10^3^ PFU ZIKV-MEX, 10^5^ PFU ZIKV-MEX, or 10^3^ PFU ZIKV-BRA on E7.5. Maternal infection was assayed by plaque assay on 2, 4, and 7 days post inoculation, and significance was determined by one-way ANOVA. (B) Rate of normal (black) versus resorbed (colored) fetuses at E11.5, E14.5, and E17.5 after maternal infection at E7.5. Data are presented as the percent of n = 23-83 total fetuses (from 3 to 10 dams per treatment group). Significance was determined by Fisher’s exact test. (C) Pregnancy outcomes of individual animals in each treatment group. Data are presented as percent of fetuses resorbed in each pregnancy. (D) Crown-to-rump length measurements in mm of morphologically normal fetuses at E11.5, E14.5, and E17.5 using ImageJ software. Significance was determined by one-way ANOVA. The color gray indicates historical data from reference (18). Significance annotations for all figures: ****, *P* ≤ 0.0001; ***, *P* ≤ 0.001; **, *P* ≤ 0.01; *, *P* ≤ 0.05.

**Table 1:** Amino acid differences between ZIKV-MEX and ZIKV-BRA. Bold text indicates deviation from other Asian-lineage ZIKVs examined in reference (Bohm 2021).

| BRA | MEX | Protein | Codon |
| --- | --- | --- | --- |
| G | **A** | NS1 | 100 |
| K | **E** | NS1 | 326 |
| **V** | M | NS1 | 349 |
| V | **I** | NS3 | 40 |
| **F** | S | NS3 | 356 |
| M | **L** | NS3 | 572 |
| **I** | **T** | NS5 | 526 |

Next, to compare the range of fetal outcomes across gestation, we necropsied dams on E11.5, E14.5, or E17.5. In an effort to minimize the use of animals, data for E14.5 for the 10^3^ PFU ZIKV-MEX and 10^3^ PFU ZIKV-BRA groups are derived from reference (18) and presented here for comparisons only. Gross examination of each conceptus revealed overt differences among fetuses within pregnancies, with uninfected counterparts, and across gestation. Fetuses appeared as either morphologically normal or undergoing embryo resorption, as defined in reference (19). The proportion of resorbed fetuses for 10^3^ PFU ZIKV-MEX, 10^5^ PFU ZIKV-MEX, and 10^3^ PFU ZIKV-BRA-infected animals did not significantly differ from PBS-inoculated controls at E11.5 (Fisher’s exact test, p > 0.1338)(**Figure 1B**). At E14.5, dams infected with 10^5^ PFU ZIKV-MEX exhibited significant fetal resorption compared to PBS-inoculated controls and 10^3^ PFU ZIKV-MEX (Fisher’s exact test, p < 0.0004) and this rate of resorption was similar to the rate caused by 10^3^ PFU ZIKV-BRA in reference (18)(**Figure 1B**). The proportion of resorbed fetuses at E14.5 for 10^5^ PFU ZIKV-MEX and 10^3^ PFU ZIKV-BRA groups was also significantly higher than what was observed at E11.5 (Fisher’s exact test, p < 0.0001)(**Figure 1B**), indicating that fetal resorption becomes grossly detectable between E11.5 and E14.5. At E17.5, the closest point to term that can be assessed in our model, the proportion of resorbed fetuses in 10^3^ PFU ZIKV-MEX-infected animals remained no different from our PBS control group (Fisher’s exact test, p > 0.0856)(**Figure 1B**), demonstrating that infection with 10^3^ PFU ZIKV-MEX does not result in significant fetal resorption at any point across gestation. Infection with 10^3^ PFU ZIKV-BRA, on the other hand, had high rates of fetal resorption at E14.5 and E17.5 that were significantly higher than PBS at all points assessed (Fisher’s exact test, p < 0.0128) but were no different from each other (Fisher’s exact test, p = 0.0875)(**Figure 1B**). The rate of fetal resorption varied significantly between individual pregnancies within each treatment group. Most groups had modest variation, but 10^5^ PFU ZIKV-MEX and 10^3^ PFU ZIKV-BRA displayed high variability at E14.5 and E17.5, ranging between 9 - 100%(**Figure 1C**).

We measured crown-to-rump length (CRL) at E11.5 and E17.5 to assess the impacts of ZIKV infection on fetal growth across gestation (18, 19, 34). Only fetuses that appeared morphologically normal were included for CRL measurement to examine intrauterine growth restriction (IUGR). There was no statistically significant difference in mean CRL in 10^3^ PFU ZIKV-MEX or 10^3^ PFU ZIKV-BRA fetuses compared to fetuses from PBS-inoculated controls at E11.5 (One-way ANOVA with Tukey’s multiple comparisons, p >0.9797) (**Figure 1D**). For 10^5^ PFU ZIKV-MEX fetuses, there was not a statistically significant reduction in CRL at E14.5 (Tukey’s multiple comparisons, p = 0.2096), which is consistent with 10^3^ PFU ZIKV-MEX fetuses but different from 10^3^ PFU ZIKV-BRA fetuses at E14.5 reported in reference (18). We observed a significant reduction in mean CRL in both 10^3^ PFU ZIKV-BRA and 10^3^ PFU ZIKV-MEX fetuses compared to PBS controls at E17.5 (One-way ANOVA with Tukey’s multiple comparisons, p <0.0001, average difference 3.24mm and 6.23mm, corresponding to an 11% and 21% reduction in fetal size, respectively). Overall, these data indicate that 10^3^ PFU ZIKV-MEX and 10^3^ PFU ZIKV-BRA both have the capacity to cause IUGR, but 10^3^ PFU ZIKV-BRA-induced IUGR manifests earlier in gestation and results in a greater magnitude of restriction.

To understand how or if infectious ZIKV virions reach the developing embryo during gestation, we examined a subset of MFI tissues for the presence of infectious virus using plaque assays. The MFI is composed of the maternal-derived decidua and the fetal-derived placenta. Consistent with our previous work (18), no infectious virus, except for one sample at E11.5, was detected by plaque assay in any fetus sample for any treatment group (**Figure 2A**). In contrast, infectious virus was detected in about one third of MFI samples from ZIKV-infected groups. At E11.5, 10^5^ PFU ZIKV-MEX MFIs had a significantly higher titer than 10^3^ PFU ZIKV-MEX MFIs (Tukey’s multiple comparisons, p = 0.0028)(**Figure 2A**). However, at E14.5, 10^3^ PFU ZIKV-BRA MFIs had a significantly higher titer than both ZIKV-MEX-inoculated groups (Tukey’s multiple comparisons, p < 0.0001)(**Figure 2A**).

**Figure 2:**
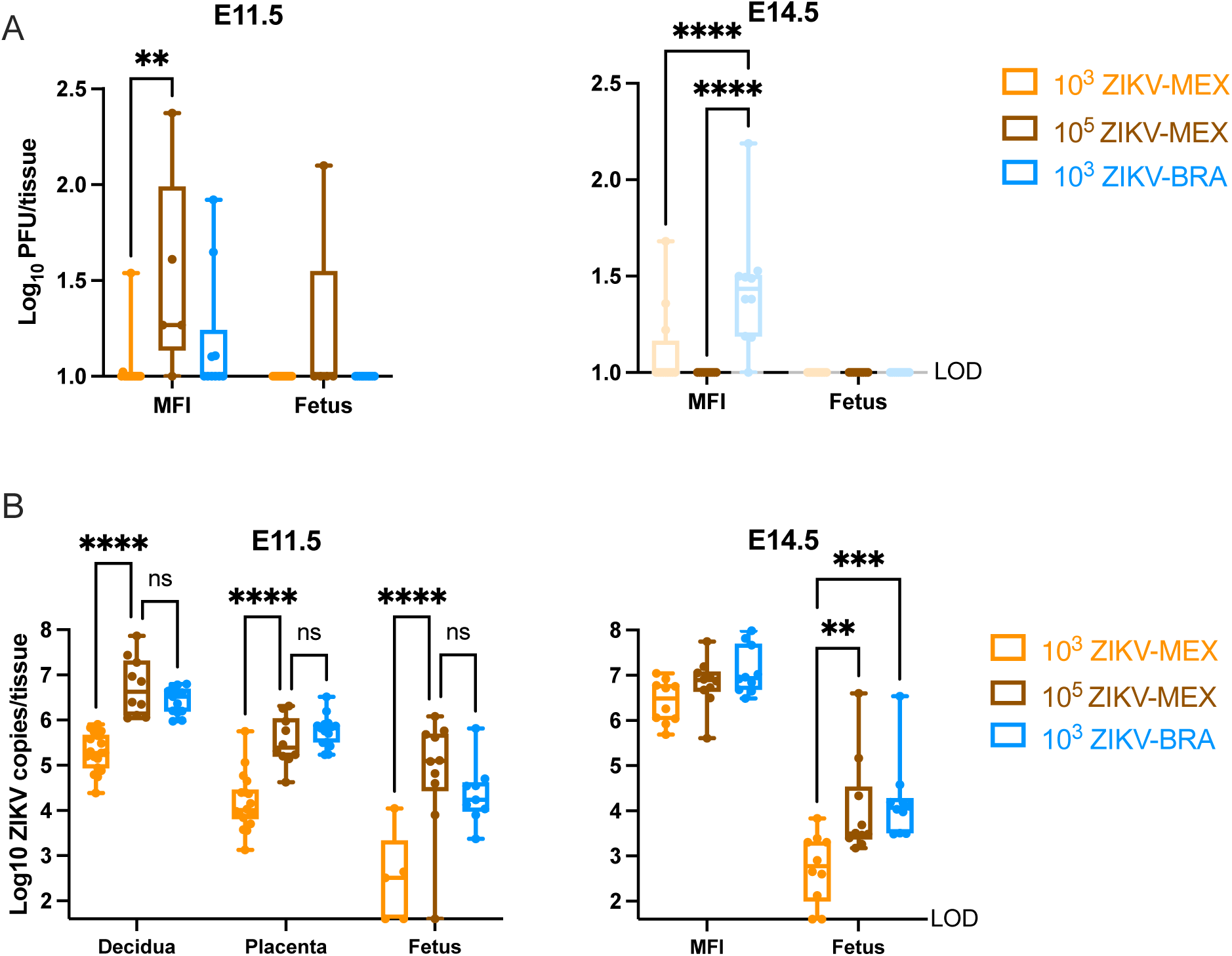
Infectious virus and ZIKV vRNA load at E11.5 and E14.5. (A) Tissue titer was measured by plaque assay for homogenized MFI (comprising decidua and placental tissues) and fetuses at E11.5 and E14.5. (B) ZIKV vRNA load was measured by qRT-PCR for homogenized decidua, placenta, MFI, and fetuses at E11.5 and E14.5. For all figures, symbols represent individual MFI or fetus samples from 4 to 10 independent experiments for each treatment group. The color gray indicates historical data from reference (Bohm 2021). Bars represent the median viral titer of each treatment group and significance was determined by two-way ANOVA with Tukey’s multiple comparisons. Significance annotations: ****, *P* ≤ 0.0001; ***, *P* ≤ 0.001; **, *P* ≤ 0.01; *, *P* ≤ 0.05; ns, *P* > 0.05.

Given the limited evidence for viral replication in fetuses or at the MFI, we next used RT-qPCR to examine these tissues for the presence of ZIKV viral RNA (vRNA)—RT-qPCR detects viable, partial, and non-viable RNA fragments. The presence of vRNA has been shown to induce antiviral signaling and synthesis of viral proteins (35), and therefore can trigger an antiviral response. We first analyzed archived MFI and fetus samples at E14.5 from reference (18). We observed no difference in MFI vRNA load between any ZIKV-inoculated groups (2-way ANOVA with Tukey’s multiple comparisons, p > 0.2470)(**Figure 2B**). We did, however, observe significantly higher fetal vRNA loads in 10^3^ PFU ZIKV-BRA and 10^5^ PFU ZIKV-MEX groups compared to 10^3^ PFU ZIKV-MEX (Tukey’s multiple comparison, p < 0.0022)(**Figure 2B**), suggesting that ZIKV-MEX vRNA can reach the fetus at the same rate as ZIKV-BRA vRNA at higher doses. Given these differences, we dissected the MFI into the maternal-derived decidua and the fetal-derived placenta at E11.5 to better understand the vRNA burden in distinct MFI structures before fetal resorption is clearly evident. At E11.5, we observed high vRNA loads in all ZIKV-inoculated groups. 10^3^ PFU ZIKV-BRA had significantly higher vRNA loads in all tissues compared to 10^3^ PFU ZIKV-MEX (p < 0.0001), but not 10^5^ PFU ZIKV-MEX at E11.5 (Tukey’s multiple comparisons, p = 0.6605). These data demonstrate that vRNA load is dependent on the dose and the genotype of the infecting ZIKV strain, with significant differences in vRNA loads observed in the decidua, placenta, and fetus prior to (E11.5) and when (E14.5) fetal resorption is detectable (**Figure 2B**).

### ZIKV influences the MFI transcriptome in a strain- and dose-dependent manner

Phenotypic characterization across gestation established that infection with 10^3^ PFU ZIKV-BRA and 10^5^ PFU ZIKV-MEX results in significantly greater fetal demise and vRNA load in MFI tissues compared to infection with 10^3^ PFU ZIKV-MEX. We therefore sought to determine how infiltration of ZIKV vRNA impacts the function of the MFI, with the aim of identifying potential mechanisms of fetal resorption. We collected deciduas and placentas from dams (n=5 per treatment group) that were inoculated with 10^3^ PFU ZIKV-MEX, 10^5^ PFU ZIKV-MEX, and 10^3^ PFU ZIKV-BRA or PBS. Decidua and placenta tissue samples were collected at E9.5 and E11.5. These timepoints were chosen because fetal resorption can be a multi-day, four-stage process (36). We therefore aimed to capture early responses that may be important for driving the resorption process. Additionally, the MFI can be dissected into functionally distinct tissues (decidua and placenta) that are large enough to isolate total RNA from a single sample without pooling. We included equal proportions of male and female decidua and placenta tissues, with one or two tissues per embryo sex per animal to avoid sex biases in our dataset. These numbers also ensured robust sampling from each pregnancy, which is critical given the broad range in fetal resorption we observed at E14.5 (see **Figure 1C**). We used DESeq2(37) to identify significantly differentially expressed genes (≥ 1 log_2_ fold, p < 0.05), Hallmark Gene Set Enrichment Analysis (Hallmark GSEA)(38, 39) to identify enriched gene families, and Pathview (40) to map differentially expressed genes to KEGG signaling pathways.

At E9.5, only five transcripts were significantly differentially expressed between PBS, 10^3^ PFU ZIKV-MEX, and 10^3^ PFU ZIKV-BRA deciduas. In contrast, 52 transcripts were significantly differentially expressed in the placenta (**Figure 3A-C**). The majority of these transcripts were differentially expressed between ZIKV-infected and PBS groups (**Figure 3A-B**), and only four transcripts differentially expressed between 10^3^ PFU ZIKV-MEX and 10^3^ PFU ZIKV-BRA (**Figure 3C**)(**Table 2**). Hallmark GSEA revealed that 10^3^ PFU ZIKV-MEX and 10^3^ PFU ZIKV-BRA E9.5 placentas were enriched for IFN alpha and gamma responses compared to PBS (**Figure 3D-E**). Hallmark gene sets are coherently expressed signatures derived by aggregating many Molecular Signature Database (MSigDB) mouse gene sets to represent well-defined biological states or processes (38, 39). The Hallmark “IFN alpha response” comprises type I and type III IFN responses. Hallmark GSEA revealed that 10^3^ PFU ZIKV-BRA E9.5 placentas are enriched for IFN alpha and gamma responses compared to 10^3^ PFU ZIKV-MEX (**Figure 3F**). Additional signatures enriched in 10^3^ PFU ZIKV-BRA compared to 10^3^ PFU ZIKV-MEX include IL-6 JAK STAT3 signaling and heme, bile acid, and xenobiotic metabolism. 10^3^ PFU ZIKV-MEX was enriched for MYC targets V1 and G2M checkpoint (**Figure 3F**). At E9.5, there was no significant fetal resorption, nor infectious virus detected in the MFI across inoculated strains and doses (**Figure 3G-H**). There were no significant differences in ZIKV vRNA loads in the decidua, placenta, or fetus samples between 10^3^ PFU ZIKV-MEX and 10^3^ PFU ZIKV-BRA (Two-way ANOVA with Sidak’s multiple comparisons, p > 0.1445)(**Figure 3I**).

**Figure 3:**
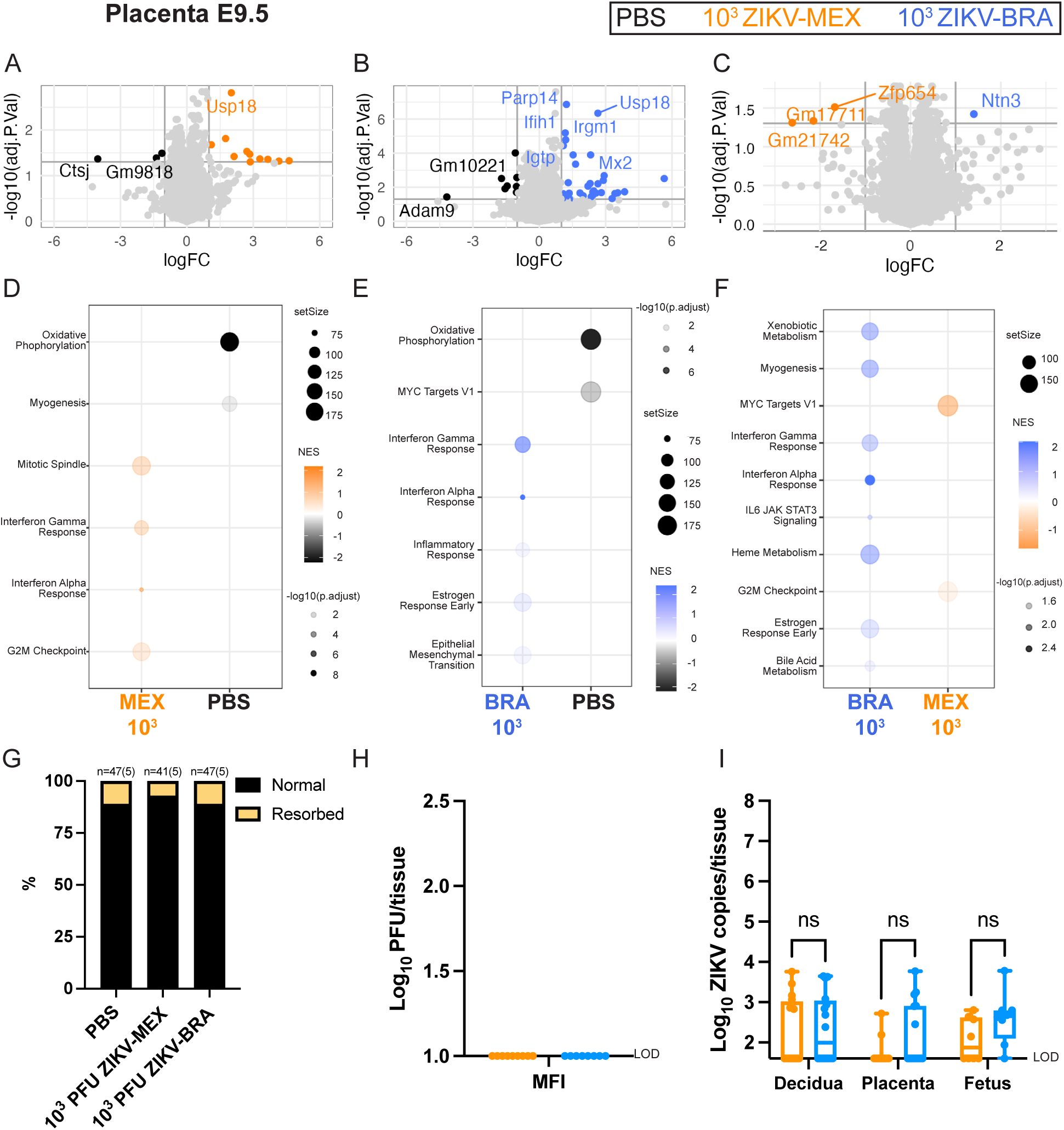
ZIKV-induced transcriptome differences in the E9.5 placenta. (A-C) Volcano plots depicting differentially expressed gene transcripts in the placenta at E9.5 of animals inoculated with 10^3^ PFU ZIKV-MEX, 10^3^ PFU ZIKV-BRA, or PBS. Genes with significant changes |log 2 fold change| >1 and -log10(p.adjust) > 0.05 appear in color; genes outside these parameters appear in light gray. (D-F) Hallmark gene set enrichment analysis of differentially expressed genes between ZIKV-infected and PBS groups. Transcriptomic data represent 16-20 embryo sex-balanced placentas from n=5 dams per inoculation group. PBS = black, 10^3^ PFU ZIKV-MEX = orange, 10^3^ PFU ZIKV-BRA = blue. (G) Rate of normal (black) versus resorbed (yellow) fetuses at E9.5 after maternal inoculation at E7.5. Data are presented as the percent of n = 41-47 total fetuses (from 5 dams per treatment group). (H) Tissue titer was measured by plaque assay for homogenized MFI (comprising decidua and placental tissues) at E9.5 for 8-9 replicates per treatment group. (I) ZIKV vRNA load in decidua, placenta, and fetuses at E9.5 was measured by qRT-PCR for 8-16 replicates per treatment group. Significance was determined by two-way ANOVA with Sidak’s multiple comparisons. Significance annotations: ns, *P* > 0.05.

**Table 2:** Differential gene expression between ZIKV-BRA and ZIKV-MEX in E9.5 placentas.

| Gene ID | Log<sub>2</sub> fold change<br>(10<sup>3</sup> PFU ZIKV-BRA vs<br>10<sup>3</sup> PFU ZIKV-MEX) | adj p<br>value | Predicted function |
| --- | --- | --- | --- |
| Ntn3 | 1.40 | 0.04 | Animal organ morphogenesis; neuron projection development;<br>and tissue development |
| Zfp654 | 1.67 | 0.03 | DNA-binding transcription factor activity, RNA polymerase II-specific, expressed in early conceptus |
| Gm17711 | 2.15 | 0.05 | Not annotated |
| Gm21742 | 2.62 | 0.05 | Not annotated |

At E11.5, we identified 179 gene transcripts that were significantly differentially expressed in the decidua, with most occurring between ZIKV-infected and PBS groups (**Figure 4A-C**). Hallmark GSEA revealed that 10^3^ PFU ZIKV-MEX, 10^5^ PFU ZIKV-MEX, and 10^3^ PFU ZIKV-BRA transcriptomes were enriched for the IFN alpha and gamma responses, as well as allograft rejection (**Figure 4D-F**). We identified multiple transcripts that were significantly differentially expressed between ZIKV-infected groups (**Figure 4G-I**). We identified transcripts that were differentially expressed based on inoculation dose (10^3^ PFU vs 10^5^ PFU)(**Figure 4G**), the inoculating ZIKV strain (ZIKV-MEX vs ZIKV-BRA)(**Figure 4H**), and between two inoculations that cause similar rates of fetal resorption (10^5^ PFU ZIKV-MEX, and 10^3^ PFU ZIKV-BRA)(**Figure 4I**). Hallmark GSEA showed that the E11.5 10^5^ PFU ZIKV-MEX decidua was enriched for the IFN alpha and gamma responses compared to 10^3^ PFU ZIKV-MEX, which was enriched for oxidative phosphorylation, G2M checkpoint, and E2F targets (**Figure 4J**). 10^3^ PFU ZIKV-BRA was also enriched for the IFN responses compared to 10^3^ PFU ZIKV-MEX (**Figure 4K**). However, when 10^5^ PFU ZIKV-MEX and 10^3^ PFU ZIKV-BRA were compared, 10^5^ PFU ZIKV-MEX was only enriched for IFN gamma and myogenesis gene sets (**Figure 4L**), indicating that these groups had similar enrichment for the type I IFN response. These data suggest that inoculum boluses containing strains or doses (10^5^ PFU ZIKV-MEX and 10^3^ PFU ZIKV-BRA) that result in significant rates of fetal demise also induce robust type I IFN responses in the decidua at timepoints just prior to fetal resorption becoming visibly detectable.

**Figure 4:**
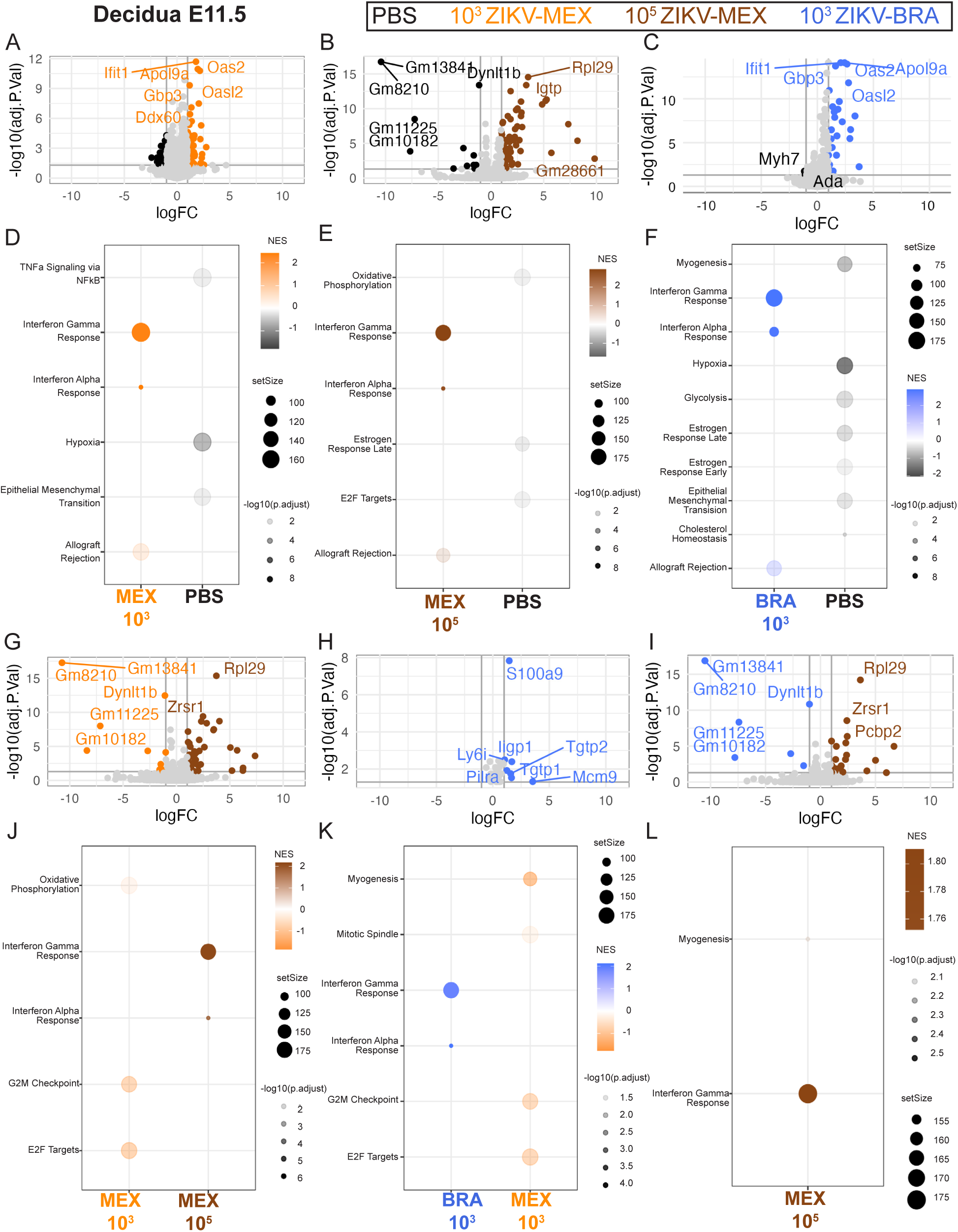
ZIKV strain and dose significantly influences the decidua transcriptome at E11.5. (A-C) Volcano plots depicting differentially expressed gene transcripts in the decidua at E11.5 of animals inoculated with 10^3^ PFU ZIKV-MEX, 10^5^ PFU ZIKV-MEX, or 10^3^ PFU ZIKV-BRA and PBS. Genes with significant changes |log 2 fold change| >1 and -log10(p.adjust) > 0.05 appear in color; genes outside these parameters appear in light gray. (D-F) Hallmark gene set enrichment analysis of differentially expressed genes between ZIKV-infected and PBS groups. (G-I) Volcano plots depicting differentially expressed gene transcripts between ZIKV-infected animals in the decidua at E11.5. Genes with significant changes |log 2 fold change| >1 and -log10(p.adjust) > 0.05 appear in color; genes outside these parameters appear in lightgray. (J-L) Hallmark gene set enrichment analysis of differentially expressed genes between ZIKV-infected E11.5 deciduas. In all figures, 10^3^ PFU ZIKV-MEX, 10^3^ PFU ZIKV-BRA, PBS data represent 14-20 embryo sex-balanced deciduas from n=4-5 dams per inoculation group. 10^5^ PFU ZIKV-MEX data represent three embryo sex-balanced deciduas from n=3 dams. PBS = black, 10^3^ PFU ZIKV-MEX = orange, 10^5^ PFU ZIKV-MEX = brown, 10^3^ PFU ZIKV-BRA = blue.

In the placenta, we identified 540 gene transcripts that were significantly differentially expressed at E11.5. Most of these differences occurred between ZIKV-infected and PBS groups (**Figure 5A-C**). Similar to our observations in the decidua, 10^3^ PFU ZIKV-MEX, 10^5^ PFU ZIKV-MEX, and 10^3^ PFU ZIKV-BRA E11.5 placentas were enriched for IFN responses and allograft rejection compared to PBS (**Figure 5D-F**). However, we also observed enrichment for the inflammatory response and MYC targets V1, suggesting that the placenta is subjected to more robust antiviral responses than the decidua.

**Figure 5:**
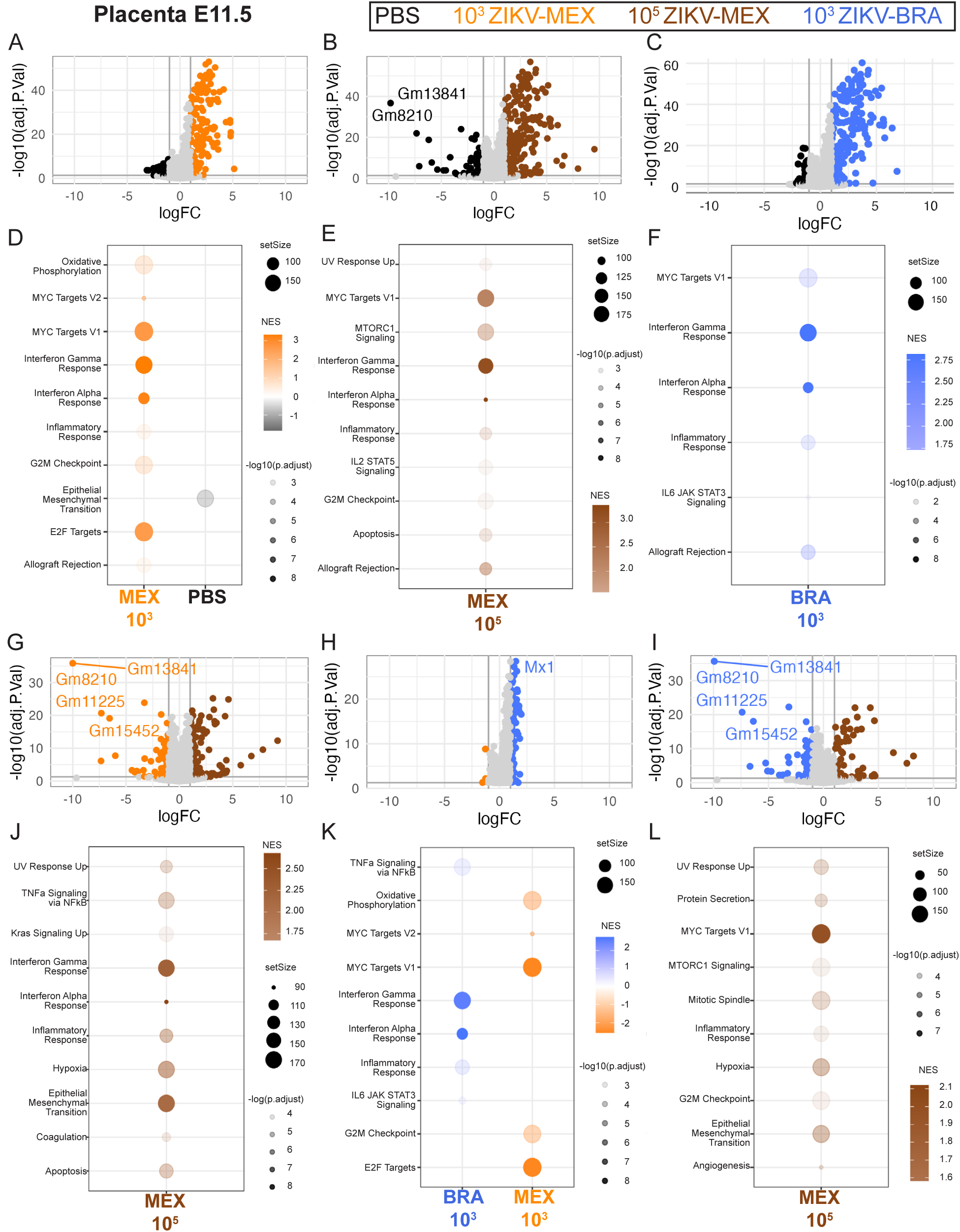
ZIKV strain and dose significantly influence the placenta transcriptome at E11.5. (A-C) Volcano plots depicting differentially expressed gene transcripts in the placenta at E11.5 of animals inoculated with 10^3^ PFU ZIKV-MEX, 10^5^ PFU ZIKV-MEX, or 10^3^ PFU ZIKV-BRA and PBS. Genes with significant changes |log 2 fold change| >1 and -log10(p.adjust) > 0.05 appear in color; genes outside these parameters appear in lightgray. (D-F) Hallmark gene set enrichment analysis of differentially expressed genes between ZIKV-infected and PBS groups. (G-I) Volcano plots depicting differentially expressed gene transcripts between ZIKV-infected animals in the placenta at E11.5. Genes with significant changes |log 2 fold change| >1 and -log10(p.adjust) > 0.05 appear in color; genes outside these parameters appear in lightgray. (J-L) Hallmark gene set enrichment analysis of differentially expressed genes between ZIKV-infected E11.5 placentas. In all figures, data represent 12-20 embryo sex-balanced placentas from n=4-5 dams per inoculation group. PBS = black, 10^3^ PFU ZIKV-MEX = orange, 10^5^ PFU ZIKV-MEX = brown, 10^3^ PFU ZIKV-BRA = blue.

We identified multiple transcripts from E11.5 placental tissue that were significantly differentially expressed among ZIKV-infected groups (**Figure 5G-I**). We identified transcripts that were differentially expressed based on inoculation dose (10^3^ PFU vs 10^5^ PFU)(**Figure 5G**), the inoculating ZIKV strain (ZIKV-MEX vs ZIKV-BRA)(**Figure 5H**), and between two inoculations that cause similar rates of fetal resorption (10^5^ PFU ZIKV-MEX, and 10^3^ PFU ZIKV-BRA)(**Figure 5I**). Hallmark GSEA revealed that the 10^5^ PFU ZIKV-MEX and 10^3^ PFU ZIKV-BRA placentas were enriched for the IFN alpha and gamma responses, inflammatory response, and TNFa signaling via NFkB compared to 10^3^ PFU ZIKV-MEX (**Figure 5J-K**). However, when 10^5^ PFU ZIKV-MEX and 10^3^ PFU ZIKV-BRA were compared, 10^5^ PFU ZIKV-MEX was enriched for the inflammatory gene set, but not TNFa signaling via NFkB nor the IFN alpha and gamma responses (**Figure 5L**), suggesting that 10^5^ PFU ZIKV-MEX and 10^3^ PFU ZIKV-BRA similarly induce these responses. Further, the enrichment scores (NES), number of genes enriched within a gene set (setSize), and adjusted p-values (-log10(p.adjust)) suggest that the IFN alpha and gamma responses are more robust than TNFa signaling via NFkB.

We mapped the placenta transcriptome at E11.5 of 10^3^ PFU ZIKV-MEX, 10^5^ PFU ZIKV-MEX, and 10^3^ PFU ZIKV-BRA (compared to PBS), to KEGG pathways using Pathview to identify homologous pathways that could be implicated in initiating the IFN response (**Figure 6**)(40). 10^5^ PFU ZIKV-MEX and 10^3^ PFU ZIKV-BRA had significant, uniform upregulation of genes in the TLR pathways, notably TLR3, which senses dsRNA (**Figure 6A-B**). When we compared 10^5^ PFU ZIKV-MEX and 10^3^ PFU ZIKV-BRA directly (**Figure 6C**), we observed variable expression of genes in TLR pathways, suggesting that these pathways were not uniformly expressed in animals that received ZIKV boluses that cause significant fetal resorption.

**Figure 6:**
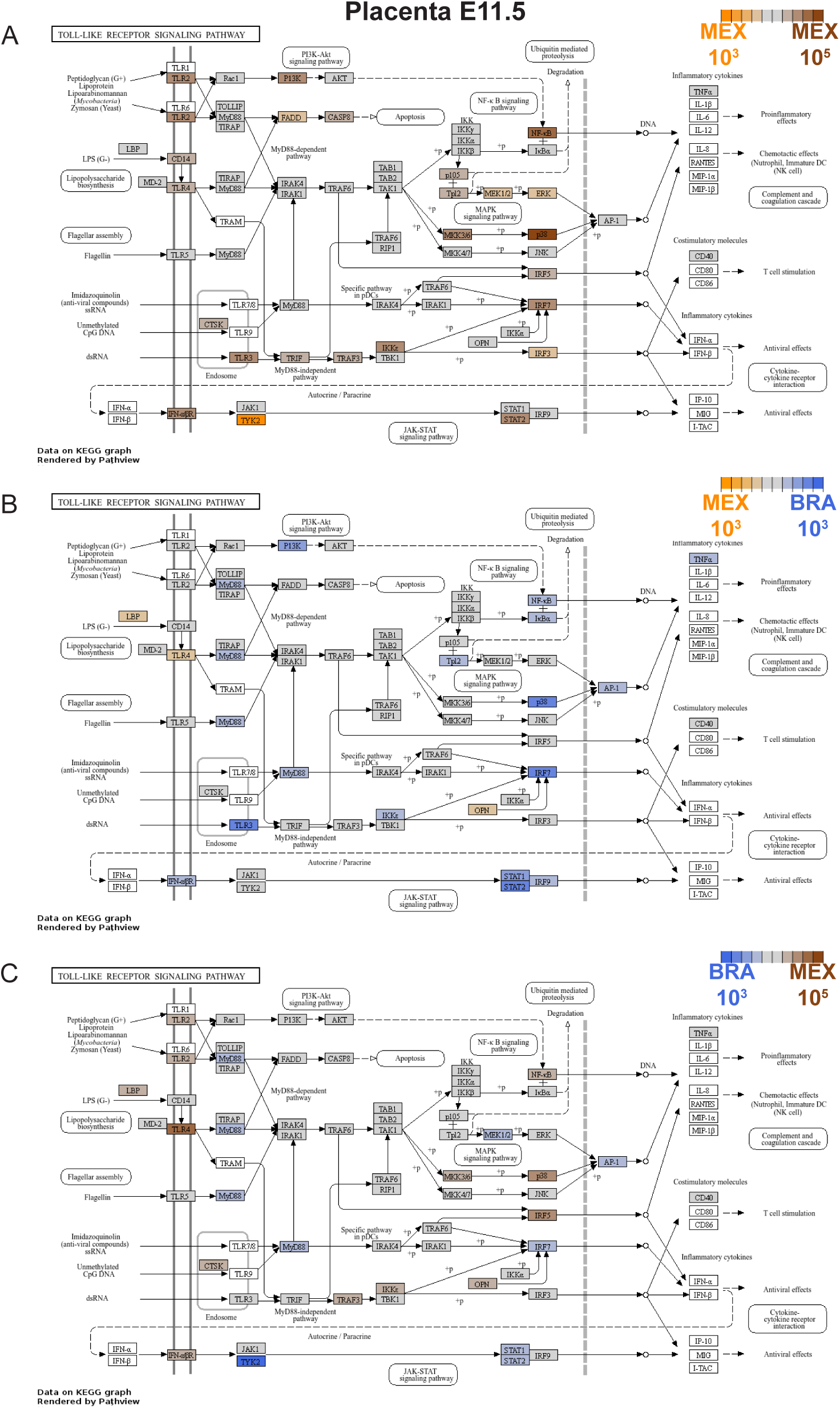
Differential expression of genes involved in the toll-like receptor (TLR) pathway by ZIKV-infected animals in the E11.5 placenta. (A-C) Genes with significant differential expression (-log10(p.adjust) > 0.05) in the E11.5 placenta were mapped to the TLR pathway using Pathview: 10^3^ PFU ZIKV-MEX = orange, 10^5^ PFU ZIKV-MEX = brown, 10^3^ PFU ZIKV-BRA = blue. Genes that were not significantly differentially expressed appear in gray. Genes not analyzed appear in white. In all figures, data represent 12-20 embryo sex-balanced placentas from n=4-5 dams per inoculation group.

When we mapped 10^5^ PFU ZIKV-MEX and 10^3^ PFU ZIKV-BRA to the RIG-I-like receptor (RLR) pathway, we observed significant, uniform upregulation of genes compared to 10^3^ PFU ZIKV-MEX (**Figure 7A-B**). In contrast to our findings with the TLR pathway, we observed almost no significant differential expression of genes in the RLR pathway between 10^5^ PFU ZIKV-MEX and 10^3^ PFU ZIKV-BRA (**Figure 7C**), suggesting that 10^5^ PFU ZIKV-MEX and 10^3^ PFU ZIKV-BRA uniformly induce RLR signaling compared to 10^3^ PFU ZIKV-MEX.

**Figure 7:**
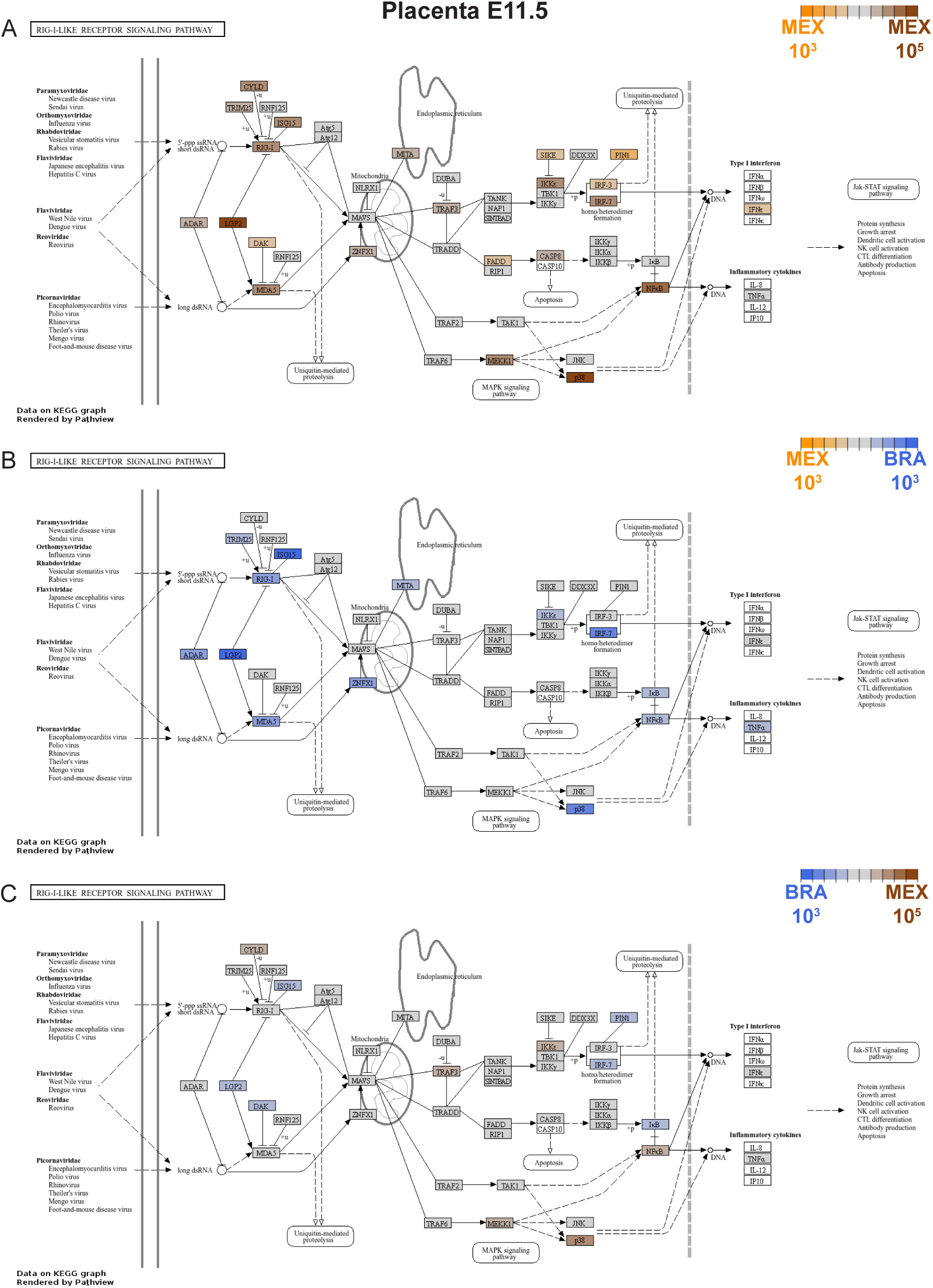
10^5^ PFU ZIKV-MEX and 10^3^ PFU ZIKV-BRA uniformly induce expression of genes in the RIG-I-like receptor (RLR) pathway in the E11.5 placenta. (A-C) Genes with significant differential expression (-log10(p.adjust) > 0.05) in the E11.5 placenta were mapped to the RLR pathway using Pathview: 10^3^ PFU ZIKV-MEX = orange, 10^5^ PFU ZIKV-MEX = brown, 10^3^ PFU ZIKV-BRA = blue. Genes that were not significantly differentially expressed appear in gray. Genes not analyzed appear in white. In all figures, data represent 12-20 embryo sex-balanced placentas from n=4-5 dams per inoculation group.

Next, we aimed to understand the association between ZIKV vRNA and the RLR-driven IFN response. ZIKV infection produces single- and double-stranded RNA intermediates during viral replication that can signal through RLRs and toll-like receptors (TLRs), which operate to induce antiviral factors including IFN and proinflammatory cytokines. We therefore plotted gene expression of IFN-response genes (*Rsad2*, *Mx1*, and *Stat2*), ZIKV pattern-recognition receptors (*Ddx58* aka RIG-I, *Ifih1* aka MDA5, and *Tlr3*), a proinflammatory cytokine (*Il1a*), and *Actin* (as a control)(**Figure 8**). We found that IFN-response genes were significantly, and proportionally expressed when compared to the vRNA load (p < 0.0001)(**Figure 8A-C**). RLR genes *Ddx58* and *Ifih1* were also significantly proportionally expressed (p <0.0005) while *Tlr3* was not (p = 0.211), suggesting that ZIKV infection preferentially induces RLR expression over other pattern-recognition receptors (**Figure 8D-F**). These results are consistent with those from our Pathview analysis, demonstrating that RLR pathways are uniformly upregulated by the two infectious boluses that result in high vRNA loads at the MFI and cause significant fetal resorption, while TLRs are not. The gene *Il1a* was not significantly proportionally expressed (p = 0.238) in relation to vRNA load suggesting that genes involved in proinflammatory cytokine production are not proportional to ZIKV vRNA (**Figure 8G**).

**Figure 8:**
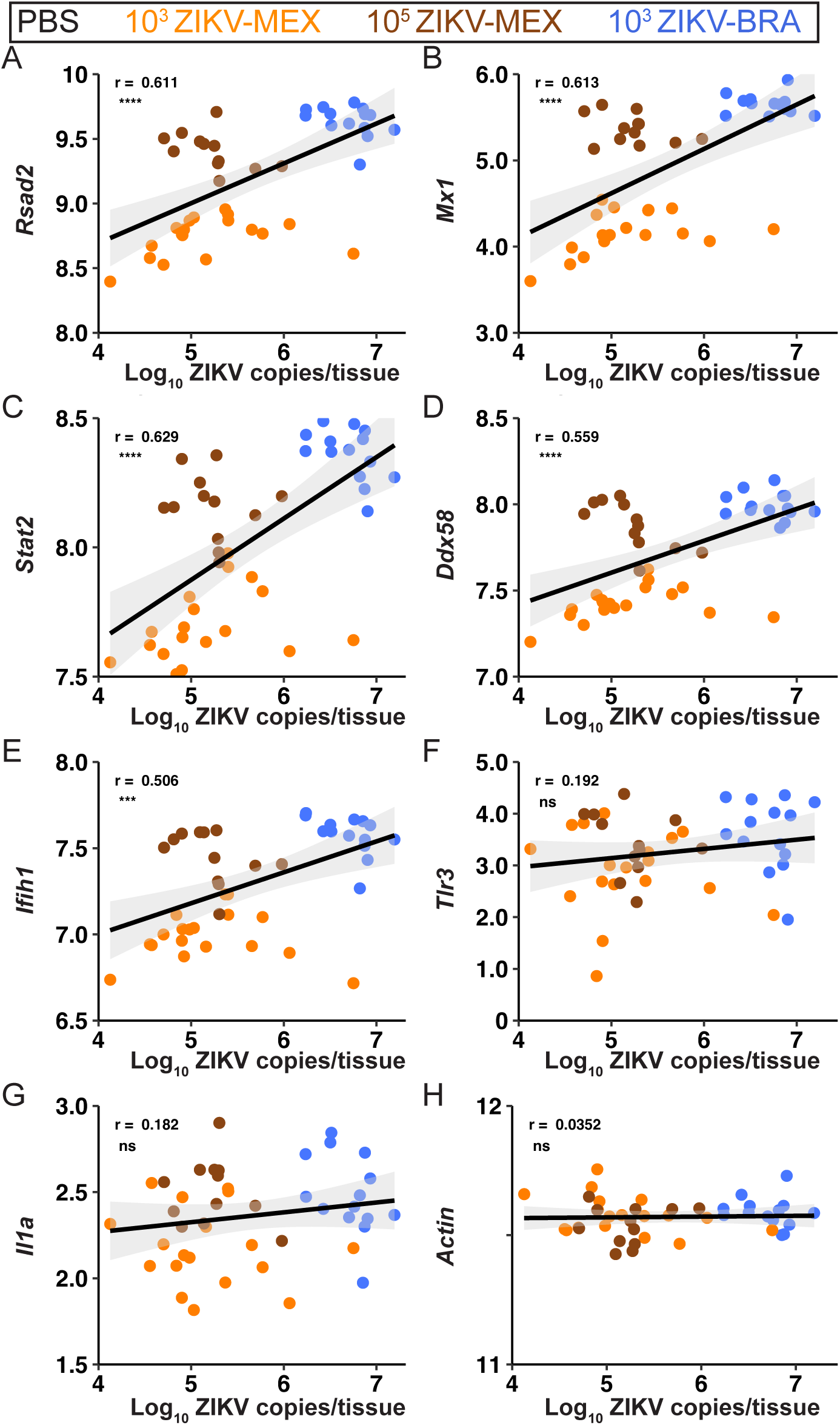
ZIKV vRNA load positively correlates with interferon-stimulated genes and RLRs. Pearson correlations with 95% confidence intervals are shown for ZIKV vRNA copies/tissue versus transcript counts (in counts per million reads) for interferon-stimulated genes *Rsad2*, *Mx1*, *Stat2* (A-C), RLR genes *Ddx58* and *Ifih1* (D-E), *Tlr3* (F), proinflammatory cytokine *Ilra* (G), and *Actin* (H). Symbols represent individual placentas from 4-5 dams inoculated with 10^3^ PFU ZIKV-MEX (orange), 10^5^ PFU ZIKV-MEX (brown), or 10^3^ PFU ZIKV-BRA (blue). Correlation coefficients (r) are shown in each panel. Significance annotations for all figures: ****, *P* ≤ 0.0001; ***, *P* ≤ 0.001; **, *P* ≤ 0.01; *, *P* ≤ 0.05; ns, > 0.05.

### Modest chemical inhibition of RIG-I activity in the placenta does not reduce the likelihood of fetal demise during ZIKV infection

The previous analyses suggested that ZIKV vRNA induces a proportional IFN response via RLRs at the MFI that can instigate fetal demise. We therefore hypothesized that vRNA sensing via RIG-I is contributing to fetal demise, because RIG-I has previously been shown to be the primary sensor of ZIKV vRNA (41, 42). To investigate this, we used RIG012, a potent chemical inhibitor of RIG-I, to reduce RIG-I activity in pregnant *Ifnar1^-/-^* mice (**Figure 9A**). RIG012 is transient in serum, but stable in tissue (**Figure 9B-C**). We therefore aimed to maximize RIG012 concentration at the MFI over the course of our experiment. We intraperitoneally injected 22.5mg/kg RIG012 every 12 hours from E6.5 - E14.5, which resulted in significant tissue permanence, averaging 0.65µM at the MFI (**Table 3**)(**Figure 9C**). This dose was well-tolerated with no signs of toxicity. A concentration of 0.65µM RIG012 is estimated to reduce RIG-I activity by ∼40% according to *in vitro* data (43). We could not dose animals with concentrations higher than this because 45mg/kg RIG012 caused lethal toxicity within 36 hours.

**Figure 9:**
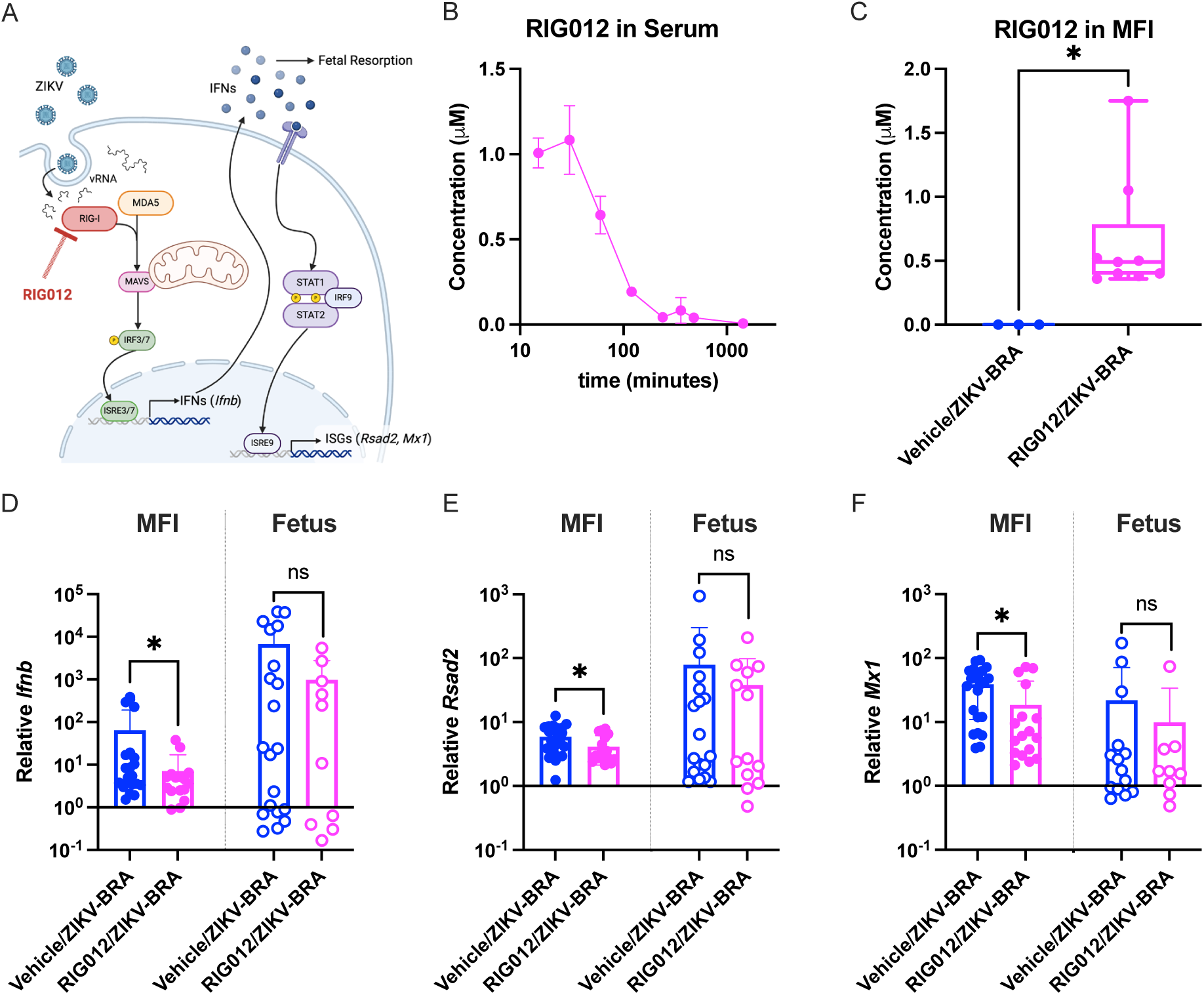
RIG012 treatment significantly inhibits RIG-I activity in the MFI. (A) Schematic of where RIG012 inhibits activity in the RLR-signaling pathway and downstream interferon (IFN) and interferon-stimulated genes (ISGs) that are expressed. (B) Concentration of RIG012 in serum of nonpregnant female mice (n=3) intraperitoneally injected once with 10mg/kg, measured by mass spectrometer. The mean with standard deviation is plotted. (C) Concentration of RIG012 in MFI at E14.5 of 10^3^ PFU ZIKV-BRA-infected animals, intraperitoneally injected every 12 hours with Vehicle or 22.5mg/kg RIG012 E6.5-E14. Whole tissue samples were homogenized in water and concentration was measured via mass spectrometer. Bars represent the median concentration and significance was determined using an unpaired t-test. Transcript abundance of *Ifnb* (D), *Rsad2* (E), and *Mx1* (F) was analyzed from MFI and fetus samples collected on E14.5 by qPCR. Expression levels were normalized to *Hprt* and the ddC*_T_* was calculated relative to samples harvested from PBS-inoculated controls. Data points represent individual samples. The mean with standard deviation is plotted. Significance was calculated with a t-test with Welch’s correction. Significance annotations for all figures: ****, *P* ≤ 0.0001; ***, *P* ≤ 0.001; **, *P* ≤ 0.01; *, *P* ≤ 0.05; ns, > 0.05.

**Table 3:** RIG012 Treatment and ZIKV-BRA infection of pregnant Ifnar1^-/-^ mice.

| Group<br>(number of animals) | Inoculation at E7.5 | Treatment |
| --- | --- | --- |
| Vehicle/PBS<br>(n=10) | PBS | Vehicle: 15uL/g DMSO/Tween80/PBS every 12 hours<br>E6.5 - E14 |
| RIG012/PBS<br>(n=7) | PBS | 22.5mg/kg RIG012 every 12 hours<br>E6.5 - E14 |
| Vehicle/ZIKV-BRA<br>(n=11) | 10<sup>3</sup> PFU ZIKV-BRA | Vehicle: 15uL/g DMSO/Tween80/PBS every 12 hours<br>E6.5 - E14 |
| Vehicle/ZIKV-BRA<br>(n=10) | 10<sup>3</sup> PFU ZIKV-BRA | 22.5mg/kg RIG012 every 12 hours<br>E6.5 - E14 |

To assess whether our RIG012 treatment schedule was sufficient to interfere with RIG-I activation *in vivo* we measured relative transcript abundance of *Ifnb*, *Rsad2*, and *Mx1* in the MFI because these genes are known indicators of RIG-I activity (43). At E14.5, the MFI of animals treated with RIG012 and challenged with 10^3^ PFU ZIKV-BRA (Vehicle/ZIKV-BRA) had significantly lower *Ifnb*, *Rsad2*, and *Mx1* expression than animals mock-treated with vehicle and challenged with 10^3^ PFU ZIKV-BRA (RIG012/ZIKV-BRA) (p = 0.049, 0.026, and 0.022, respectively)(**Figure 9D-F**). Fetuses from Vehicle/ZIKV-BRA and RIG012/ZIKV-BRA groups had no difference in relative *Ifnb*, *Rsad2*, and *Mx1* expression (p = 0.057, p = 0.463, and p = 0.437, respectively).

To evaluate the extent to which RIG012 treatment protects against ZIKV-induced fetal demise, we subcutaneously inoculated RIG012-treated and vehicle-treated pregnant *Ifnar1^-/-^* mice in the footpad with 1 × 10^3^ PFU ZIKV-BRA, or phosphate buffered saline (PBS) to serve as experimental controls. The proportion of resorbed fetuses for RIG012/PBS did not differ significantly from Vehicle/PBS (16% vs. 11%; Fisher’s exact test, p = 0.4083)(**Figure 10A**). Consistent to what we have reported previously (18), Vehicle/ZIKV-BRA induced a rate of resorption that was significantly higher than Vehicle/PBS group (43% vs. 11%; Fisher’s exact test, p < 0.0001)(**Figure 10A**). However, no differences were observed in the proportion of resorbed fetuses in RIG012/ZIKV-BRA groups compared to Vehicle/ZIKV-BRA groups (41% vs 43%; Fisher’s exact test, p = 0.8861)(**Figure 10A**), demonstrating that at this dose, RIG012 treatment did not protect from ZIKV-induced fetal demise.

**Figure 10:**
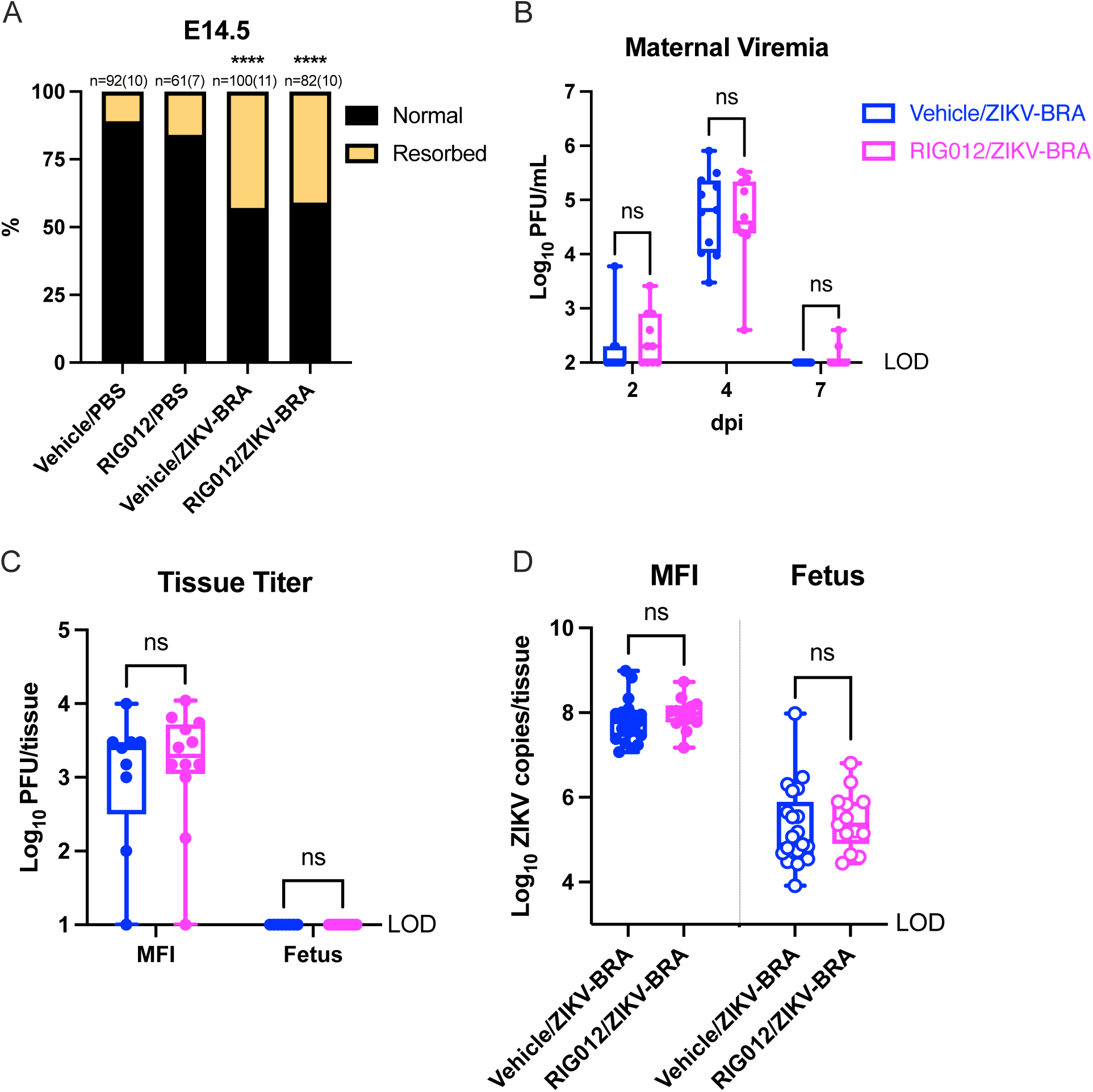
RIG012 treatment does not protect against fetal demise. (A) Time-mated *Ifnar1*^−/−^ dams were treated with Vehicle or 22.5mg/kg RIG012 every 12 hours from E6.5-E14, inoculated with 10^3^ PFU ZIKV-BRA on E7.5, and the rate of resorption was calculated at E14.5. Data are presented as the percent of n = 61-100 total fetuses (from 7 to 10 dams per treatment group). Significance was determined by Fisher’s exact test. (B) Maternal viremia was assessed via plaque assay at 2, 4, and 7 days post inoculation (dpi) and significance was determined by two-way ANOVA with Tukey’s multiple comparisons. (C) Tissue titer was assessed via plaque assay of MFI and fetus samples harvested at E14.5 and significance was determined by two-way ANOVA with Sidak’s multiple comparisons (D) ZIKV vRNA load was assessed via qRT-PCR of MFI and fetus samples harvested at E14.5. Significance was determined by unpaired t-test. For all figures: ****, *P* ≤ 0.0001; ***, *P* ≤ 0.001; **, *P* ≤ 0.01; *, *P* ≤ 0.05; ns, > 0.05.

We collected serum at 2, 4, and 7 dpi to compare viremia kinetics between Vehicle- and RIG012-treated animals. There were no significant differences in serum titers between Vehicle/ZIKV-BRA and RIG012/ZIKV-BRA at any time point (Two-way ANOVA, p > 0.9999) (**Figure 10B**). At E14.5 we collected MFI and fetal tissues; we used plaque assay to quantify infectious virus present and qRT-PCR to determine ZIKV vRNA loads. We found no significant difference in infectious virus at the MFI, and fetuses had undetectable levels of infectious virus (Two-way ANOVA with Sidak’s multiple comparisons, p > 0.9990)(**Figure 10C**). There was no significant difference in ZIKV vRNA load of the MFI and fetus between RIG012- and Vehicle-treated animals (t-test with Welch’s correction, p = 0.3218 and p = 0.5515, respectively)(**Figure 10D**).

To evaluate whether there were differences in the MFI vRNA load or IFN response based on fetal outcome, we compared the vRNA load and relative transcript abundance of *Ifnb*, *Rsad2*, and *Mx1* between normal and resorbed concepti (“Outcome”) among Vehicle/ZIKV-BRA and RIG012/ZIKV-BRA (“Treatment”) groups (**Figure 11A-D**). Unexpectedly, the vRNA load of the MFI did not significantly differ between normal and resorbed Outcomes (Two-way ANOVA with Tukey’s multiple comparisons, p = 0.106). None of the genes we compared were significantly differentially expressed in response to an interaction between Treatment and Outcome (Two-way ANOVA with Tukey’s multiple comparisons, p > 0.241). Fetal Outcome was not significantly associated with MFI expression of *Ifnb*, *Rsad2*, and *Mx1* (Two-way ANOVA with Tukey’s multiple comparisons, p > 0.291)(**Figure 11A-D**).

**Figure 11:**
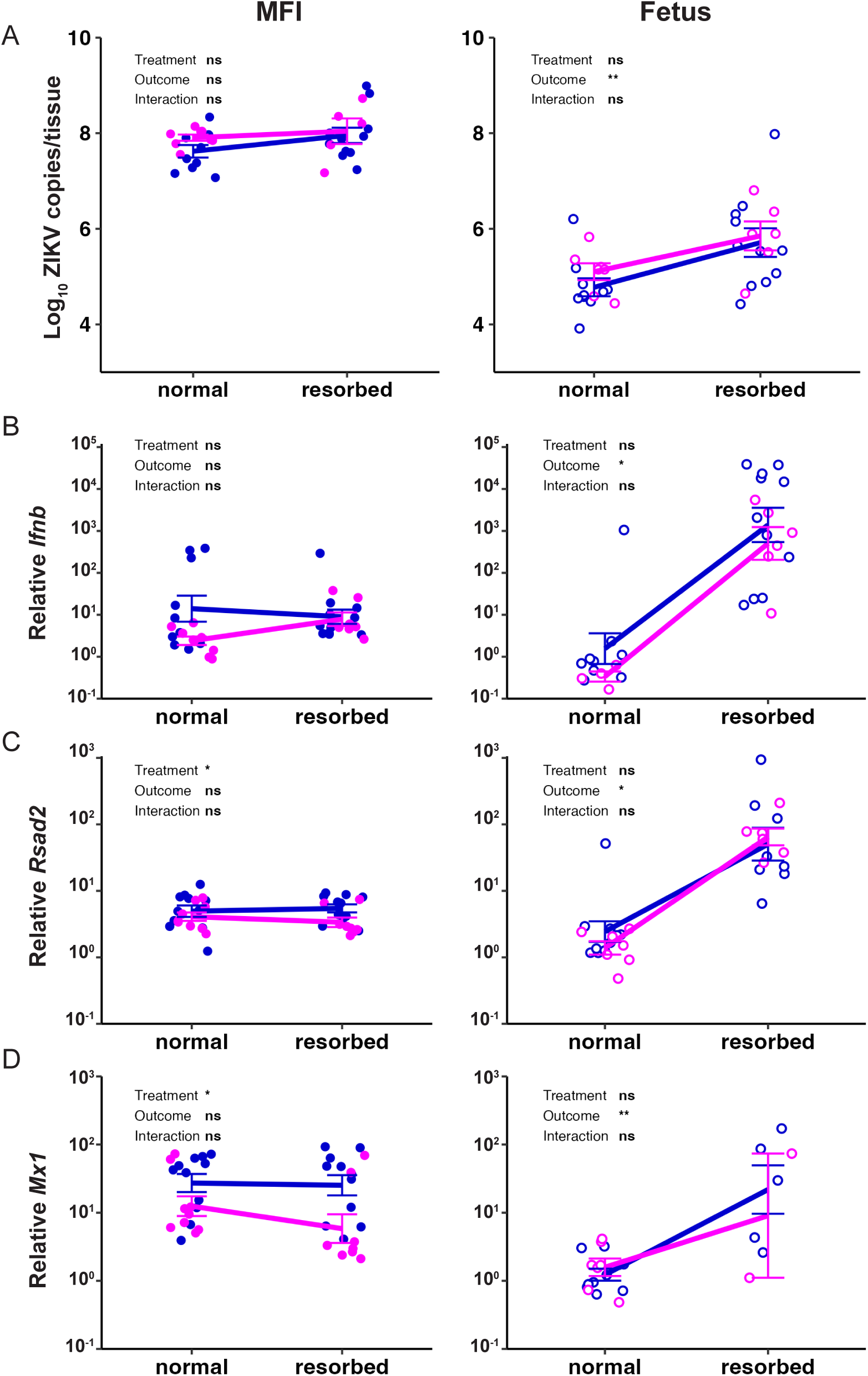
Resorbed fetuses have significantly higher relative interferon-stimulated gene expression than their normal counterparts. ZIKV vRNA load (A), relative *Ifnb* (B), *Rsad2* (C), and *Mx1* (D) expression in the MFI and fetus were plotted against Outcome (resorbed vs normal fetal outcome), and separated by Treatment (Vehicle vs 22.5mg/kg RIG012). Two-way ANOVA with Tukey’s multiple comparisons was used to determine significance. For all figures: ****, *P* ≤ 0.0001; ***, *P* ≤ 0.001; **, *P* ≤ 0.01; *, *P* ≤ 0.05; ns, > 0.05.

In contrast, we observed a significant association between fetal Outcome, fetal vRNA load, and fetal relative *Ifnb*, *Rsad2*, and *Mx1* expression (Two-way ANOVA with Tukey’s multiple comparisons. p < 0.041)(**Figure 11A-D**). Resorbed fetuses had, on average, 1 log_10_ ZIKV copies/tissue more than normal fetuses (Two-way ANOVA with Tukey’s multiple comparisons, p = 0.002). Resorbed fetuses also had nearly 10^4^ higher relative *Ifnb* abundance (Two-way ANOVA with Tukey’s multiple comparisons, p = 0.019), 10^2^ higher relative *Rsad2* abundance (Two-way ANOVA with Tukey’s multiple comparisons, p = 0.041), and 10^1.7^ higher relative *Mx1* abundance (Two-way ANOVA with Tukey’s multiple comparisons, p = 0.005), compared to normal fetuses. Overall, resorbed fetuses had higher vRNA loads and IFN-stimulated gene expression than their normal counterparts.

## DISCUSSION

Here, we expanded on our previous work (18), to demonstrate that ZIKV strain-dependent phenotypic heterogeneity is driven by antiviral immune signaling at the MFI and/or fetus. These observations substantially contribute to our nascent understanding of the mechanisms by which ZIKV harms the developing fetus. Our finding, that ZIKV activates a robust IFN response in the MFI prior to fetal resorption, is consistent with observations from other studies that mostly support a role for hyperinflammatory and/or hyperimmune responses as mediators of adverse fetal outcomes during congenital viral infections (17, 21, 44–47). For example, experiments using a breeding scheme that enabled the examination of pregnant dams that carry a mixture of fetuses that express type I IFN signaling (*Ifnar1^+/-^*) or do not express type I IFN signaling (*Ifnar1^-/-^*) within the same uterus found that only *Ifnar1^+/-^* fetuses were resorbed after ZIKV infection during early pregnancy, whereas their *Ifnar1^-/-^*littermates continued to develop (17). Similarly, experiments using mice lacking the IFN lambda (IFN-λ) receptor found that IFN-λ can have either a protective antiviral effect or cause immune-mediated pathology, depending on the stage of gestation when IFN-λ signaling occurs (21). Interestingly, the protective and pathogenic effects of IFN-λ occurred through signaling in maternal immune cells rather than in fetal or placental tissues. In contrast, and in the setting of maternal immunocompetence, mitochondrial antiviral-signaling protein (MAVS)-dependent type I IFN signaling in the fetus was found to be necessary to restrict ZIKV infection in the fetal compartment of the placenta (48). Here we observe ZIKV strain- and dose-dependent RLR-mediated activation of the IFN response at the MFI and identify a significant fetal IFN response that correlates with fetal resorption.

When the ZIKV genome is replicated in the cytoplasm of a host cell it produces multiple ssRNA and dsRNA intermediates. These ZIKV vRNAs are primarily recognized by RIG-I, which recognizes the 5’ region of the ZIKV genome, which triggers the production of type I IFN and proinflammatory cytokines (41, 42). RNA binding triggers a confirmation change in RIG-I that promotes interaction with MAVS. Viral sensing via RIG-I and downstream signaling via MAVS are transiently induced by the host to restrict viral replication (48, 49). However, if vRNA persists, the host is inundated with an aberrant RIG-I-driven IFN response and this prolonged RIG-I signalling can trigger immunopathology (50–52). We had therefore posited that prolonged RIG-I sensing of ZIKV vRNA may be an important driver of adverse pregnancy outcomes during ZIKV infections, possibly due to increased type I IFN production (53–55). Indeed, our results showed significant enrichment for IFN responses in the decidua and placenta prior to significant fetal resorption, with positive correlation between ZIKV vRNA load and IFN-stimulated genes, but not *Tlr3* or the proinflammatory cytokine *Il1a*. However, chemical inhibition of RIG-I in the MFI via RIG012 treatment had no effect on the rate of fetal resorption following inoculation with 10^3^ PFU ZIKV-BRA, suggesting that inhibition of RIG-I signalling in the MFI is not sufficient to protect the feto-placental unit—at least at the doses tested here. Critically, results may have differed had we been able to achieve more robust inhibition of RIG-I signalling. However, this was not possible because of RIG012-associated toxicity at higher doses. We chose not to investigate this phenomenon in RIG-I knockout mice because complete ablation of RIG-I sensing could result in uncontrolled viral replication in the dam, thus failing to recapitulate the specific mechanism of fetal harm observed herein. A possible useful alternative could involve using breeding schemes involving *Ddx58*^-/-^ mice (note that the *Ddx58* gene encodes murine *Rig-i*) crossed with *Ifnar1*^+/+^, *Ifnar1*^+/-^, and *Ifnar1*^-/-^ mice. This may help better disentangle the role of RLR-driven immunopathology at the MFI and subsequent fetal demise, however it is important to note that of number of *Ddx58*^-/-^ mouse models are embryo lethal (56) or develop spontaneous colitis from commensal viruses (57–59) and therefore would not be suitable for examining pathologic outcomes following ZIKV-infection during pregnancy.

Another possible explanation for differences in fetal outcomes observed between treatment groups could be that ZIKV vRNA also binds TLRs that, in turn, activate IFN responses (60); however, recent work determined that TLR7/8, TLR9, MyD88, STING are not substantially involved in antiviral activity in the fetus and placenta (48). And, surprisingly, *MyD88*^-/-^ fetuses (downstream of TLR7/8 and TLR9) resulted in lower viral burden in the decidua and placenta than those with intact *MyD88* (48). In contrast, binding of TLR3 by ZIKV vRNA suppresses the RIG-I-driven IFN response and promotes viral replication (61). Importantly, we observed inconsistent and incomplete differential activation of TLR pathways during pathologic ZIKV infections (10^5^ PFU ZIKV-MEX and 10^3^ PFU ZIKV-BRA). We therefore maintain that RIG-I-mediated IFN activation is a more likely mediator of fetal resorption in the *Ifnar1^-/-^* model.

Because resorbed fetuses had significantly higher ZIKV vRNA loads and relative levels of the interferon-stimulated genes *Rsad1, Ifnb,* and *Mx1* compared to normal fetuses and normal and resorbed placentas, we speculate that the fetal, rather than the placental, immune response is an important driver of fetal resorption. Indeed, Fetal Inflammatory Response Syndrome is known to be caused by systemic activation of fetal IFNs and this can result in neurological complications or death (62, 63) similar to what has been observed from infections with teratogenic pathogens like ZIKV, but more studies are needed to understand the relative importance of fetal-derived immune responses. As previously mentioned, a prior study found that *Ifnar1*^-/-^ fetuses were protected from fetal resorption while *Ifnar1*^+/-^ fetuses were not (17), but the fetal IFN response was not examined so its contribution to fetal resorption in that system remains unknown. Further, in an immunocompetent mouse model, the IFN response was more robust in fetal endothelial cells compared to placental cells (48), suggesting that the magnitude of the response may determine its contribution to resorption.

While the IFN response appears to be a primary mediator of fetal demise in the *Ifnar1*^-/-^ model, it is important to consider the possibility that this phenotype is multifactorial. For example, the 10^5^ PFU ZIKV-MEX placenta transcriptome had significant enrichment for MYC targets V1, hypoxia, and epithelial mesenchymal transition compared to 10^3^ ZIKV-BRA. MYC targets V1 are associated with cell proliferation (64), suggesting that 10^5^ PFU ZIKV-MEX placentas experienced greater tissue growth compared to 10^3^ ZIKV-BRA. Because cell proliferation is closely linked with apoptosis (65), enrichment for MYC targets V1 may indicate compensation for cell death that is occurring. In fact, enrichment for MYC targets V1 was observed in all of our ZIKV-inoculated groups when compared to PBS. Hypoxia-induced changes in metabolism drive placentation in mice and humans (66), however after placentation, hypoxia conditions can increase inflammation through release of damage-associated molecular patterns (DAMPs)(67). At certain levels, inflammation and DAMPs increase the risk of intrauterine growth restriction and stillbirth, even in the absence of a pathogen(67). Murine placentation is complete at E10.5, suggesting that enrichment for hypoxia in the E11.5 placenta is detrimental (68). Enrichment for epithelial mesenchymal transition suggests a greater presence of migratory cells (69), which is critical for formation of the labyrinth and gastrulation (68). Poor labyrinth formation would impact nutrient and gas exchange between mother and fetus (70), which could result in intrauterine growth restriction and fetal death. Abnormal gastrulation would impact cell type and location during embryo development (71), which could result in an improperly formed embryo. While these signatures may be secondary to a robust IFN response induced by ZIKV-MEX and ZIKV-BRA, they have important implications for potential concurrent mechanisms of fetal resorption.

ZIKV-MEX and ZIKV-BRA are genetically very similar, but differences observed in fetal outcomes between the two strains may be due to virus genetic determinants of virulence. The seven amino acid differences between them occur in the NS1, NS3, and NS5 proteins (**Table 1**). ZIKV NS1 disrupts endothelial barrier function (72), which is particularly important at the placenta because endothelial cells remodel the maternal and fetal placental vasculature. Abnormalities in placental endothelial cells lead to high rates of apoptosis, and subsequent fetal growth restriction and pre-eclampsia (73). It is possible that ZIKV-BRA may produce higher levels of NS1 compared to ZIKV-MEX and therefore may be more adept at disrupting endothelial barriers, thus contributing to significantly higher rates of fetal resorption—but we did not test that here. This could also explain why ZIKV-MEX is capable of causing fetal demise at higher doses. ZIKV NS3 binds dsRNA replication intermediates and associates with NS5 to promote genome replication, and mutations in the ATPase or RNA-binding region of ZIKV NS3 have both been shown to alter helicase activity and reduce genome replication (74). Therefore, it is possible that differences in NS3 helicase activity between the two strains may explain the different ZIKV vRNA loads observed in the decidua, placenta, and fetus. Further, ZIKV NS3 has been associated with brain calcifications in ZIKV-infected fetuses (75), demonstrating that the overall activity and concentration of ZIKV NS3 can be associated with adverse outcomes.

Importantly, CD8 T cell epitopes are located in NS1, NS3, and NS5 (76). Therefore, polymorphisms at these sites between ZIKV-MEX and ZIKV-BRA may alter T cell activation, including differentially inducing cytotoxic CD8 T cells, but more studies are needed to investigate this. During congenital infection and/or hyperinflammatory states, maternal and fetal CD8 T cells infiltrate the MFI (77, 78). ZIKV activation of CD8 T cells has been associated with significant IFN gamma, TNF alpha, and granzyme B production (79–81), all of which are cytotoxic, despite being required to control ZIKV infection (76, 82). CD8 T cells induce cytotoxic effects in response to ZIKV in immunologically privileged spaces like the neuronal cavity (83), but their role at the MFI remains unknown. Other congenital infections, including human cytomegalovirus, induce maternal- and fetal-derived CD8 T cell-mediated cytotoxic effects in the placenta (78, 84, 85), and can even mediate allogeneic intolerance (86). In fact, one study found that ZIKV-infected placentas from fetuses with microcephaly had increased T cell activation, suggesting that T cell activation plays a role in the severity of CZS (87). Future work should consider how T cells, particularly CD8 T cells, mediate pathology during ZIKV-infected pregnancies. The future spread of ZIKV will remain a threat to pregnant people in many locations around the globe. While the exact mechanism underlying ZIKV-induced fetal harm remains unclear, these studies highlight that RIG-I can mediate a pathologic IFN response at the MFI and that the fetal immune response may be an underappreciated contributor to adverse pregnancy outcomes during ZIKV infections.

## METHODS

*Ethical approval.* This study was approved by the University of Minnesota, Twin Cities Institutional Animal Care and Use Committee (Animal Care and Use protocol number 2401-41654A).

### Cells and Viruses

African green monkey kidney cells (Vero cells; ATCC CCL-81) were maintained in Dulbecco’s modified Eagle medium (DMEM) supplemented with 10% fetal bovine serum (FBS; Corning, Manassas, VA), 1× Antibiotic Antimycotic solution (Corning, Manassas, VA) and incubated at 37°C in 5% CO2. *Aedes albopictus* mosquito cells (C6/36; ATCC CRL-1660) were maintained in DMEM supplemented with 10% fetal bovine serum (FBS; HyClone, Logan, UT), 2 mM L-glutamine, 1.5 g/liter sodium bicarbonate, 1× Antibiotic Antimycotic solution, and incubated at 28°C in 5% CO2. The cell lines were obtained from the American Type Culture Collection, were not further authenticated, and were not specifically tested for mycoplasma. ZIKV strain R116265 (ZIKV-MEX; GenBank KX766029) was originally isolated from a 73-year-old-male traveling in Mexico in 2016 with a single round of amplification on Vero cells (CDC, Ft. Collins, CO). ZIKV strain Paraiba_01 (ZIKV-BRA; GenBank KX280026) was originally isolated from human serum in Brazil in 2015 with two rounds of amplification on Vero cells, and a master stock was obtained from Kevin Noguchi at Washington University in St. Louis (St. Louis, MO). Virus challenge stocks were prepared by inoculation onto a confluent monolayer of C6/36 mosquito cells. Virus challenge stocks were sequence authenticated as described in reference (18).

### Plaque Assay

Quantification of virus titer in maternal serum, placenta, and fetuses were completed by plaque assay on Vero cells. Duplicate wells were infected with 0.1 ml aliquots from serial 10-fold dilutions in growth medium and virus was adsorbed for 1 h. After incubation, the monolayers were overlaid with 3 ml containing a 1:1 mixture of 1.2% oxoid agar and 2× DMEM (Gibco, Carlsbad, CA) with 10% (vol/vol) FBS and 2% (vol/vol) Antibiotic Antimycotic solution. Cells were incubated at 37°C in 5% CO2 for 3 days (ZIKV-BRA) or 5 days (ZIKV-MEX) for plaque development. Cell monolayers were then stained with 3 ml of overlay containing a 1:1 mixture of 1.2% oxoid agar with 4% neutral red (Gibco) and 2× DMEM with 2% (vol/vol) FBS, and 2% (vol/vol) Antibiotic Antimycotic solution. Cells were incubated overnight at 37°C in 5% CO2 and plaques were counted.

### Mice

Female *Ifnar1*^−/−^ mice on the C57BL/6 background were bred in the specific pathogen-free animal facilities of the University of Minnesota within the College of Veterinary Medicine. Male C57BL/6 mice were purchased from Jackson Laboratories. Timed matings between female *Ifnar1^−/−^* mice and male C57BL/6 mice resulted in *Ifnar1^+/−^* progeny.

### Subcutaneous Inoculation

All pregnant dams were between 6 and 10 weeks of age and were randomly assigned to infected or control groups. Matings between *Ifnar1^−/−^*dams and wild-type sires were timed by checking for the presence of a vaginal plug, indicating gestational age E0.5. At embryonic day 7.5 (E7.5) dams were inoculated in the right hind footpad with 1 × 10^3^ or 1 x 10^5^ PFU of the selected ZIKV strain in sterile phosphate-buffered saline (PBS) or with sterile PBS alone to serve as experimental controls. All animals were closely monitored by laboratory staff for adverse reactions and/or clinical signs of disease. A submandibular blood draw was performed at 2, 4, 7 and/or 10 days post inoculation (dpi), and serum was collected to verify viremia. Mice were humanely euthanized and necropsied at E9.5, E11.5, E14.5, or E17.5.

### Intraperitoneal administration of RIG012

RIG012 (MedChemExpress, Monmouth Junction, NJ) was dissolved in sterile DMSO at a concentration of 30mg/mL before being mixed with an equal volume of Tween 80 and stored at 4°C. Mice were weighed and doses were calculated. The RIG012 in DMSO/Tween 80 solution was diluted with nine parts sterile water immediately prior to injection to make a final concentration of 5/5/90 (DMSO/Tween 80/H_2_O) which was dosed at 15uL/g to provide a dose of 22.5mg/kg. A control solution of 5/5/90 (DMSO/Tween 80/H_2_O) was dosed at 15uL/g. Animals were intraperitoneally injected using a 28G needle with a 1mL syringe. Animals were monitored for signs of toxicity for up to 1 hour post injection, and every 12 hours following injection.

### Mouse necropsy

Following inoculation with ZIKV or PBS, mice were sacrificed at E9.5, E11.5, E14.5, or E17.5. Tissues were carefully dissected using sterile instruments that were changed between each mouse to minimize possible cross contamination. Each organ and neonate was morphologically evaluated *in situ* prior to removal. Using sterile instruments, the uterus was removed and dissected to remove individual concepti. Each conceptus was placed in a sterile culture dish and dissected to separate the fetus and the maternal-fetal interface (MFI) for gross evaluation. Fetuses were characterized as “normal” or “resorbed,” with the latter being defined as having significant growth retardation and reduced physiological structure compared to littermates and controls, accompanied by clearly evident developmental delay or visualization of a macroscopic plaque in the uterus. The MFI included maternal-derived decidua tissue and fetal-derived placental tissue. At E9.5 and E11.5, the MFI was further dissected under a stereoscope to separate decidua and placenta tissues. Tissues isolated at E9.5, E11.5, and E17.5 were snap frozen in RNase-free tubes on dry ice. Tissues isolated at E14.5 were snap frozen as described or frozen in PBS supplemented with 20% FBS and 1% Antibiotic Antimycotic. A subset of tissues from each timepoint were fixed in 10% neutral buffered formalin for 24 to 96 hours (depending on tissue mass) then transferred to 70% ethanol until imaged.

### Crown-to-rump length

Crown-to-rump length (CRL) was measured by tracing the distance from the crown of the head to the base of the tail, using ImageJ. Resorbed fetuses were excluded from measurement analyses because they would not survive if the pregnancy was allowed to progress to term (19).

### Fetal and MFI viral titers

An Omni TH115 homogenizer (Omni International, Kennesaw, GA) was used to homogenize fetus and MFI samples following necropsy. Samples were submerged in chilled PBS supplemented with 20% FBS and 1% Antibiotic Antimycotic solution in 2 ml Safelock tubes (Eppendorf, Hamburg, Germany). Omni soft tissue probes (Omni International, Kennesaw, GA) were used to homogenize samples at medium speed. Homogenized samples were clarified by centrifugation at 10,000 × *g* for 2 min. The supernatant was removed and 0.1 ml was immediately plated in duplicate for plaque assay. The remainder was stored at −80°C.

### Determination of fetal sex

DNA was extracted and purified from E9.5 and E11.5 fetuses using a Zymo Quick-DNA miniprep plus kit (Zymo Research, Irvine, CA) or Maxwell RSC Tissue DNA kit (Promega, Madison, WI). PCR and gel electrophoresis were conducted as previously described (88).

### Total RNA extraction

Total RNA was extracted and purified from deciduas, placentas, and fetuses using a Direct-zol RNA miniprep kit (Zymo Research, Irvine, CA). RNA was eluted in 50 to 100uL RNase-free water. RNA concentration and purity were measured by a Qubit 4 fluorometer (ThermoFisher, Waltham, MA).

### Quantification of vRNA load

Viral RNA was quantified from extracted total RNA from maternal-fetal tissues by quantitative reverse transcription-PCR as described previously (18, 19, 89). Total RNA was titrated by qRT-PCR using TaqMan Fast virus 1-step master mix (Applied Biosystems, Waltham, MA) on a QuantStudio3 (ThermoFisher, Waltham, MA). ZIKV RNA titers were interpolated from a standard curve of diluted in vitro-transcribed ZIKV RNA. The limit of detection for this assay is 150 ZIKV genome copies/ml (1.60 log10 copies/tissue).

### Illumina RNAseq library preparation and sequencing

Multiplex sequencing libraries were generated from 500 ng of total RNA (per library) using Illumina’s TruSeq sample prep kit and multiplexing sample preparation oligonucleotide kit (Illumina Inc., San Diego, CA) following the manufacturer’s instructions. Up to four samples per tissue per animal per inoculation group, with equal proportions male and female, were submitted for sequencing. Samples were sequenced on an Illumina NovaSeq, which generated 2x150 bp paired-end reads at a depth of 20 million reads. Illumina’s bcl2fastq v2.20 was used for de-multiplexing, and sequence quality was assessed based on %GC content, average base quality, and sequence duplication levels.

### Sequence alignment and transcript quantification

RNA sequencing data were quality-checked using FastQC (v0.11.9)(90) and summarized using MultiQC (v1.12)(91). The resulting trimmed reads were aligned to the *Mus musculus* genome [Mus_musculus.GRCm39.cdna.all.index] using kallisto (v0.46.1)(92), which relies on a pseudoalignment framework. Out of 3.7 billion sequence reads, 73–93% of reads mapped unambiguously to the *Mus musculus* reference genome. Downstream analysis followed the DIY Transcriptomics R workflow (93) in R (v4.2.3), supplemented by Pathview analysis (40). Aligned reads were annotated using the tximport (v1.28.0) R package (94). Differentially expressed genes were identified using raw gene counts. Differential gene expression analysis was performed using the DESeq2 package (v1.40.1)(37) using a significance cutoff of 0.05 and a fold change cutoff of 1 log_2_ fold change. Volcano plots, temporal plots, and heatmaps were generated using the ggplot2 package (v3.4.2) in R (95). Gene Set Enrichment Analysis was performed using GSEA (v4.3.2)(39) on normalized data against Hallmark gene sets available from MSigDB (Mouse MSigDB Collections 2004). All data processing and analysis scripts are publicly available on GitHub (https://github.com/aliotalab/ZIKVplacentaRNAseq/tree/main).

### Quantification of RIG012 in serum and maternal-fetal interface (MFI) tissue

5 µL plasma samples were directly loaded to a 96-well Millipore Multiscreen Solvinert 0.45 micron low binding PTFE hydrophilic filter plate. MFI samples were homogenized with water (x3 dilution) then 5 µL was loaded to the filter plate. All plasma/tissue samples were treated with 75 µL 90/10 acetonitrile/water with Atorvastatin as I.S. to extract the analyte and precipitate protein. The plates were agitated on ice for approximately ten minutes prior to centrifugation into a collection plate. Separate standard curves were prepared in blank mouse plasma and tissue homogenate and processed in parallel with the samples. The filtrate was directly analyzed by LC-MS/MS analysis against. HPLC and MS/MS parameters are provided in the accompanying tables (**Table 4 - Table 6**).

**Table 4:** LC (Shimadzu UFLC XR) conditions.

| Compound | RIG012 | I.S. (Atorvastatin) |
| --- | --- | --- |
| Column | Thermo Betasil C18 5µ, 50x2.1mm |  |
| Mobile phase | A: Water with 0.1% Formic Acid<br>B: Acetonitrile with 0.1% Formic Acid |  |
| Flow rate (ml/min) | 0.35 |  |
| Temperature (°C) | 35 |  |
| Injection volume (µl) | 10 |  |

**Table 5:** Gradient elution conditions.

| Time (min) | Mobile phase A (%) | Mobile phase B (%) |
| --- | --- | --- |
| 0.2 | 90 | 10 |
| 0.5 | 90 | 10 |
| 2.0 | 5 | 95 |
| 3.0 | 5 | 95 |
| 4.0 | 90 | 10 |
| 5.9 | 90 | 10 |

**Table 6:** MS (API6500+) conditions.

| Compound | RIG012 | I.S. (Atorvastatin ) |
| --- | --- | --- |
| MRM(-) | 359.4/268.2 | 557.1/397 |
| Collision Gas | Low |  |
| Curtain GAS | 30 |  |
| Ion Source Gas1 | 55 |  |
| Ion Source Gas2 | 55 |  |
| Ion Spray Voltage | -4500 |  |
| Temperature (°C) | 550 |  |
| Collision Energy | -26 | -50 |
| Declustering Potential | -75 | -75 |
| Entrance Potential | -10 |  |
| Collision Cell Exit Potential | -10 |  |

### Gene Expression of RIG-I-induced genes

RNA was extracted and purified from placentas using a Direct-zol RNA kit (Zymo Research). The High-Capacity RNA-to-cDNA kit (Applied Biosystems) was used to synthesize cDNA. Quantitative PCR using Fast Advanced Master Mix (TaqMan) was used to quantify RIG-I-induced genes on a QuantStudio3 (Applied Biosystems). The following TaqMan assays were used: *Hprt* (Mm00446968_m1), *Ifnb* (Mm00439552_s1), *Rsad2* (Mm00491265_m1), and *Mx1* (Mm00487796_m1). *Ifnb*, *Rsad2*, and *Mx1* were normalized to *Hprt* and then the threshold cycle value (2-delta delta C_T_) was calculated relative to Vehicle/PBS controls.

### Statistical analyses

All statistical analyses from the pathology data were conducted using GraphPad Prism 9 (GraphPad Software, CA, USA) or RStudio (Posit Software, PBC, Boston, MA, USA). Statistical analyses from the transcriptomic data were conducted in RStudio, under the null hypothesis of equal gene expression between groups. Statistical significance was designated to P-values of less than 0.05.

### Data availability

Raw Illumina sequencing data are available on the NCBI Sequence Read Archive under BioProject no. **SUB15084024** (https://www.ncbi.nlm.nih.gov/bioproject/SUB15084024). All data processing and analysis scripts are publicly available on GitHub (https://github.com/aliotalab/ZIKVplacentaRNAseq/tree/main).

## ACKNOWLEDGEMENTS

We thank the University of Minnesota’s Genomics Center for RNA sequencing and the Minnesota Supercomputing Institute for computing resources. We thank Dr. Vivian Bardwell for training on early gestation necropsies and Dr. Micah Gearhart for advising on bioinformatic analysis. We thank Dr. Grace Vaziri for her critical review of the manuscript.

